# Tar patties are hotspots of hydrocarbon turnover and nitrogen fixation during a nearshore pollution event in the oligotrophic southeastern Mediterranean Sea

**DOI:** 10.1101/2023.07.16.546273

**Authors:** Maxim Rubin-Blum, Yana Yudkovsky, Sophi Marmen, Ofrat Raveh, Alon Amrani, Ilya Kutuzov, Tamar Guy-Haim, Eyal Rahav

## Abstract

Weathered oil, that is, tar, forms hotspots of hydrocarbon degradation by complex biota in marine environment. Here, we used marker gene sequencing and metagenomics to characterize the communities of bacteria, archaea and eukaryotes that colonized tar patties and control samples (wood, plastic), collected in the littoral following an offshore spill in the warm, oligotrophic southeastern Mediterranean Sea (SEMS). We show aerobic and anaerobic hydrocarbon catabolism niches on tar interior and exterior, linking carbon, sulfur and nitrogen cycles. Alongside aromatics and larger alkanes, short-chain alkanes appear to fuel dominant populations, both the aerobic clade UBA5335 (*Macondimonas*), anaerobic Syntropharchaeales, and facultative Mycobacteriales. Most key organisms, including the hydrocarbon degraders and cyanobacteria, have the potential to fix dinitrogen, potentially alleviating the nitrogen limitation of hydrocarbon degradation in the SEMS. We highlight the complexity of these tar-associated communities, where bacteria, archaea and eukaryotes co-exist, exchanging metabolites and competing for resources and space.

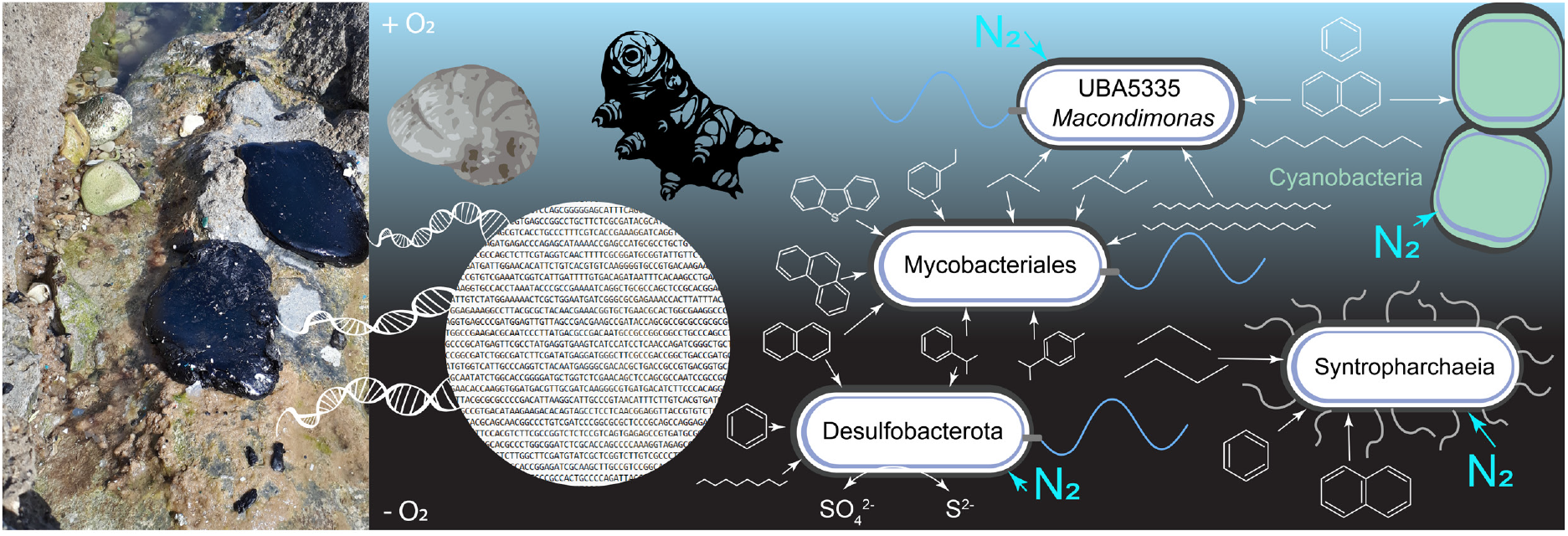

## Introduction

Oil pollution is a prominent threat to marine and coastal ecosystems, especially in the Mediterranean Sea, where petroleum transportation is substantial (Ciappa, 2023; Ezra et al., 2000; Polinov et al., 2021). Oil spills have a pronounced effect on marine life (Alford et al., 2014; Almeda et al., 2013; Brussaard et al., 2016; Gutierrez, 2018; Helm et al., 2015), altering planktonic communities (González et al., 2009; Ozhan et al., 2014; Shai et al., 2021; Tang et al., 2019), shifting microbial abundance, diversity and function (Brussaard et al., 2016; Gutierrez et al., 2013; Hazen et al., 2010; Kimes et al., 2014) and having deleterious effects on zooplankton communities (Abbriano et al., 2011; Almeda et al., 2016, 2013; Negri et al., 2016). To effectively mitigate these adverse impacts, it is crucial to understand and monitor the processes that happen before, during and after the oil pollution at the community level. Omics is a key modern tool to assess this issue, allowing robust functional modeling of the system at multiple trophic levels (Harik et al., 2022). The biodegradation of fossil fuels is catalyzed by taxonomically-diverse microbes, that cooperate and exhibit a broad range of functions and substrates, including aromatic and aliphatic hydrocarbons (Head et al., 2006). Specific bacterial lineages, including *Alcanivorax*, *Marinobacter*, *Thallassolituus*, *Cycloclasticus*, *Oleispira* and others are often detected in marine oil spills and considered obligate hydrocarbonoclastic bacteria (OHCB) (Yakimov et al., 2022, 2007). When oxygen is depleted, other organisms are associated with fossil hydrocarbons, such as short-chain alkane-degrading Syntropharchaeia (Laso-Peréz et al., 2016) and Desulfobacterota that either ferment aromatics or couple their oxidation to reduction of sulfate and iron (Ramos et al., 2014). Anaerobic and aerobic lineages synergistically degrade oil when redox gradients are present (Sierra-Garcia et al., 2020). The dynamics of the hydrocarbon-degrading bacteria throughout the pollution event are complex, whereas nitrogen limitation of oil degradation can potentially stimulate nitrogen fixation (Chronopoulou et al., 2013). Metagenomics has been widely used to provide insights into the functionality of these hydrocarbon-degrading communities in different locations and spill scenarios (Harik et al., 2022; Hazen et al., 2010; Hidalgo et al., 2020; Mason et al., 2014; Redmond and Valentine, 2012; Ribicic et al., 2018; Sierra-Garcia et al., 2020; Techtmann and Hazen, 2016).

Natural and anthropogenic liquid petroleum releases into the sea can be transformed by weathering, sedimentation, and other processes into tar balls, tar mats, and tar patties, which can accumulate nearshore and pollute the coastal and littoral environments (Warnock et al., 2015). These tar balls or patties are a prominent source of polycyclic aromatic hydrocarbons (PAHs) and n-alkanes (Elshafie et al., 2007; Wang et al., 2023; Zahari et al., 2021) that fuel specialized microbial communities (Bacosa et al., 2016; Tran et al., 2019). These microbes form biofilms on the tar surface, stimulating hydrocarbon sequestration (McGenity, 2014). For example, synthetic consortia of *Alcanivorax, Marinobacter* and *Pseudomonas* were shown to remove most of the n-alkanes and PAHs from a tarball within 45 days (Shinde et al., 2020). Tar patties and tarballs are not only hotspots of microbial activity but also provide a structural habitat (rafts) for various non-microbial organisms, including algae and metazoans such as cirripeds, which likely colonize the particles following the temporal succession paradigm (Bergmann et al., 2015; Eriksen et al., 2019; Minchin, 1996; Thiel, 2003; Thiel and Gutow, 2005). Yet, the diversity and functionality of biota associated with weathered oil are not fully understood, and, in particular, the microbial functions are not studied to the same extent as those of liquid petroleum degraders.

Here we focus on biota associated with the nearshore weathered oil pollution in the warm, oligotrophic southeastern Mediterranean Sea (SEMS). In contrast to the well-studied marine systems affected by petroleum discharge, such as the Gulf of Mexico, the SEMS is highly oligotrophic and characterized by low primary production, chlorophyll-*a* concentrations and extremely low levels of dissolved nutrients in surface waters (Ben Ezra et al., 2021; Berman-Frank and Rahav, 2012; Kress et al., 2014). Phytoplankton productivity is also limited in the SEMS coastal waters despite sporadic moderate eutrophication (Rahav et al., 2018; Rahav and Bar-Zeev, 2017; Raveh et al., 2015). Coastal water temperature ranges between 17 and 31°C throughout the year, and it is warming following the basin trends (Ozer et al., 2020; Rilov, 2016). To date, only a handful of studies attempted to better understand the response of the intrinsic microbiota to the hydrocarbon enrichment in the SEMS, mainly based on microcosm and mesocosm studies (Kababu et al., 2022; Liu et al., 2017; Shai et al., 2021).

In February 2021, the shoreline and littoral along Israel’s Mediterranean coast were severely polluted with oil in different stages of weathering, including tar balls and patties (Herut et al., 2021). The leak from a vessel occurred on February 2^nd^, whereas oil reached Israel’s shoreline on February 17^th^, being proclaimed by the authorities as a tier 2B spill. This pollution event co-occurred with a severe storm that washed large quantities of waste ashore and from the shore, polluting the littoral area not only with the weathered oil but also with plant and plastic debris. Using metagenomics and amplicon sequencing, we aimed to investigate the diversity and functions of biota associated with tar patties that were collected from nearshore tidal pools and were heterogeneously weathered. We studied the communities that formed biofilms on tar exterior, as well as those that thrive in the anaerobic interior, using tar samples that were collected from 4 days up to a month following the event. To differentiate between ambient biota, biofilm generalists and specialist tar colonizers, we also analyzed samples of tidal pool water, as well as other particles, including plastic and plant debris. We focused on prokaryotic microbes but also explored the diversity of eukaryotes.

## Materials and methods

### Sample collection

Tar samples were collected 4, 19, and 28 days following the event (on February 21^st^, March 7^th^ and March 16^th^), from the littoral tidal pools nearshore Israel (Tel Shikmona, Haifa, 32.826° N, 34.956° E), following a pollution and storm events in February 2021 (Figure S1). On February 21^st^, we collected two types of samples – partially dehydrated tar patties that were exposed to air on the rocky littoral and viscous tar patties floating in tidal pools. On March 7^th^ and March 16^th^, we collected only the viscous patties (**Figure 1**). For four tar samples, we separated the soft surface from the solidified core using a scalpel blade. Additional samples of biofilms on the water surface, small pieces of degraded plant material, and three different types of small plastic litter – sheets, filaments and white squares, were collected on March 16^th^ (**Supplementary Figure S1**). An additional sample for chemical analyses was collected on February 21^st^ (**Supplementary Note 1**).

**Figure 1:**
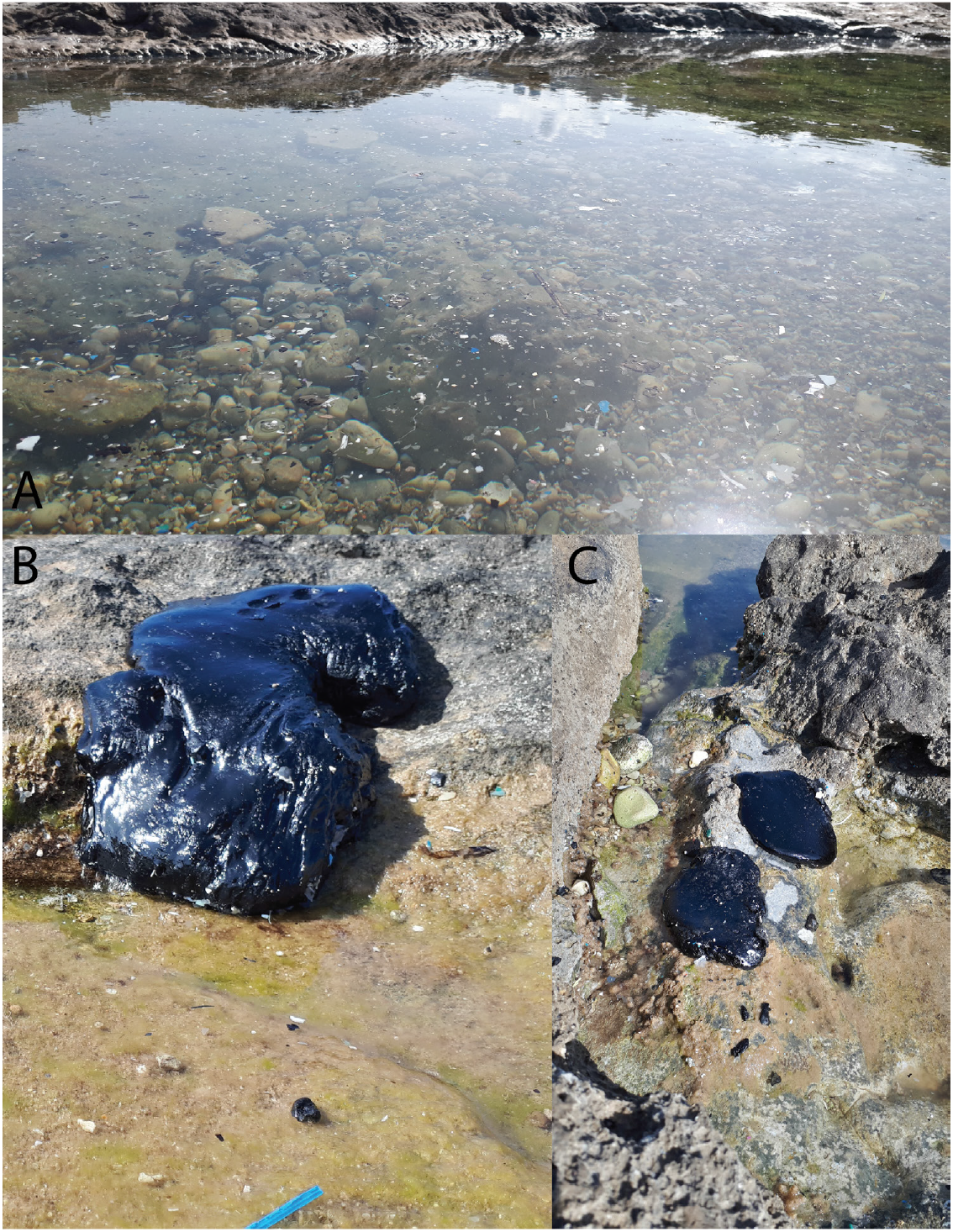
Littoral tar pollution (Tel Shikmona, Haifa, Israel, images taken on March 16, two weeks following the pollution event, samples C-F). A) Tar and other debris (plastic, plant organics) in a tidal pool following a pollution event and a storm. B,C) Large tar patties (10-30 cm) washed out to the rocky Tel Shikmona beach.

### DNA extraction and sequencing

DNA was extracted from 16 tar samples (February 21^st^ – 8, March 7^th^ – 4, March 16^th^ – 4), as well as plant organics (1 sample) and plastics (3 samples), using the legacy DNeasy PowerSoil Kit (Qiagen) (Table 1). Tidal pool water (∼200 ml) and water with white biofilm (∼10 ml) were filtered on Supor membranes (diameter 15 mm, pore size 0.2 μm, Pall). The membranes were cut into small pieces in a sterile petri dish, and the DNA was extracted using the same kit, using the FastPrep-24™ Classic (MP Biomedicals, USA) bead-beating to disrupt the cells (2 cycles at 5.5 m sec^-1^, with a 5 min interval). The V4 region (∼ 300 bp) of the 16S rRNA gene was amplified from the DNA (∼50 ng) using the 515Fc/806Rc primers amended with CS1/CS2 tags (5’-ACACTGACGACATGGTTCTACAGTGYCAGCMGCCGCGGTAA, 5’-TACGGTAGCAGAGACTTGGTCTGGACTACNVGGGTWTCTAAT, (Apprill et al., 2015; Parada et al., 2016). PCR conditions were as follows: initial denaturation at 94 °C for 45 s, 30 cycles of denaturation (94 °C for 15 sec), annealing (15 cycles at 50 °C and 15 cycles at 60 °C for 20 sec) and extension (72 °C for 30 s). Two annealing temperatures were used to account for the melting temperature of both forward (58.5-65.5 °C), and reverse (46.9-54.5 °C), primers. Library preparation from the PCR products and sequencing of 2x250 bp Illumina MiSeq reads was performed by HyLabs (Israel). For metagenomics, 10 libraries were constructed, whereas DNA from several samples with similar communities determined by the amplicon sequencing was mixed (**Table 1**). The samples were selected to cover large taxonomic diversity of tar-associated microbes. Metagenomic libraries were constructed by HyLabs, Israel, and sequenced as circa 70 million 150 bp paired-end reads using the Illumina HiSeq at GENEWIZ (Germany).

**Table 1:** The list of samples analyzed in this study. The letters in parentheses identify the tar patties, e.g., samples 1 and 2 originate from tar patty A. NA - Metagenome not sequenced.

| Specimen | Collection Date | Description | Metagenomic library |
| --- | --- | --- | --- |
| 1 | 21.02.21 | Viscous tar from the tidal pool, inner hardened (A) | T69 |
| 2 | 21.02.21 | Viscous tar from the tidal pool, outer soft (A) | NA |
| 3 | 21.02.21 | Viscous tar from the tidal pool, inner hardened (B) | T70 |
| 4 | 21.02.21 | Viscous tar from the tidal pool, outer soft (B) | T72 |
| 5 | 21.02.21 | Viscous tar from the tidal pool, inner hardened (C) | T70 |
| 6 | 21.02.21 | Viscous tar from the tidal pool, outer soft (C) | T72 |
| 7 | 21.02.21 | Dehydrated tar from the littoral rock (D) | T71 |
| 8 | 21.02.21 | Dehydrated tar from the littoral rock (E) | T71 |
| 9 | 07.03.21 | Viscous tar from the tidal pool, outer soft, biofilm (F) | T24 |
| 10 | 07.03.21 | Viscous tar from the tidal pool, outer soft (G) | T25 |
| 11 | 07.03.21 | Viscous tar from the tidal pool, outer soft (G) | NA |
| 12 | 07.03.21 | Viscous tar from the tidal pool, outer soft (H) | NA |
| 13 | 16.03.21 | Viscous tar from the tidal pool, outer soft, visible biofilm (F) | T20 |
| 14 | 16.03.21 | Viscous tar from the tidal pool, inner soft, below biofilm (F) | T21 |
| 15 | 16.03.21 | Viscous tar from the tidal pool, outer soft, visible biofilm (G) | NA |
| 16 | 16.03.21 | Viscous tar from the tidal pool, tiny pieces (H) | NA |
| 17 | 16.03.21 | Tidal pool water | NA |
| 18 | 16.03.21 | Tidal pool water with white biofilm | NA |
| 19 | 16.03.21 | Degraded plant material | T22 |
| 20 | 16.03.21 | Plastic debris – white squares | T23 |
| 21 | 16.03.21 | Plastic debris – filaments | T23 |
| 22 | 16.03.21 | Plastic debris – sheets | T23 |

### Bioinformatics

For the 16S rRNA gene amplicons, the demultiplexed paired-end reads were processed in QIIME2 V2022.2 environment (Bolyen et al., 2019). Primers and sequences that didn’t contain primers were removed with cutadapt (Martin, 2011), as implemented in QIIME2. Reads were truncated based on quality plots, checked for chimeras, merged and grouped into amplicon sequence variants (ASVs) with DADA2 (Callahan et al., 2016). The amplicons were classified with scikit-learn classifier that was trained either on Silva database V138 (16S rRNA, Glöckner et al., 2017). Mitochondrial and chloroplast sequences were removed from the dataset. Downstream analyses were performed in R V3.6.3 (R Core Team, 2023), using packages phyloseq (McMurdie and Holmes, 2013), ampvis2 (Andersen et al., 2018). Differential abundance was tested using DESeq2 (Love et al., 2014).

Metagenomic libraries were processed using ATLAS V2.0 (Kieser et al., 2020), with SPAdes V3.14 de-novo (--meta) assembler (Prjibelski et al., 2020) following read merging and error correction with the BBtools suite (Bushnell, B., https://sourceforge.net/projects/bbmap/), metaBAT2 (Kang et al., 2015), maxbin2 (Wu et al., 2015) and VAMB (Nissen et al., 2021) as binners, and DAS Tool (Sieber et al., 2018) as a final binner. We used CheckM2 to assess the completeness of the metagenome-assembled genomes (MAGs) (Chkolovski et al., 2022; Parks et al., 2015). We assigned MAG taxonomy with GTDB-tk V1.15 (Chaumeil et al., 2020) and annotated them with the SEED and the rapid annotation of microbial genomes using Subsystems Technology (RAST) (Overbeek et al., 2014), DRAM V1.4 (Shaffer et al., 2020) and METABOLIC V4 (Zhou et al., 2022). RemeDB (Sankara Subramanian et al., 2020) and CANT-HYD (Khot et al., 2022) databases and their integrated tools were used to analyze the metabolic potential for the degradation of hydrocarbons. Mechanisms of extracellular electron transfer in MAGs were identified with FeGenie (Garber et al., 2020), while the multiheme cytochromes MHCs were identified by searching ORFs for ≥3 CxxCH, CxxxCH, CxxxxCH, CxxCK and A/FxxCH motifs and prediction of localization using PSORTb V3.0.3 (Li et al., 2023; Yu et al., 2010).

The phylogenies of AcrA/McrA, PhmoC and SmoX proteins were constructed with MEGA v11 (Tamura et al., 2021) or FastTree (Price et al., 2009), following sequence alignment with MAFFT (Katoh and Standley, 2013). The phylogenomic tree based on the GTDB-tk treeing of the Genome Taxonomy Database marker gene set alignment was visualized with iTOL version 5 (Letunic and Bork, 2021).

### Nitrogen fixation rates

Surface seawater samples were collected on April 3rd, 2022 in the offshore SEMS (2 m depth, 32.95 N, 34.46 E). The seawater was pre-filtered onto 100 µm mesh to remove large-size mesozooplankton into six 4.6 L transparent Nalgene bottles. Three bottles were amended with tar patties collected in February 2021 (∼0.5 g L-1), alongside unamended controls. The ^15^N_2_-enriched seawater was prepared before the cruise by dissolving ^15^N_2_ gas (Cambridge Isotopes, lot #NLM-363-PK) in degassed seawater at a 1:100 ratio (vol: vol) (Mohr et al., 2010). We added 5% ^15^N_2_-enriched seawater to the incubation containers after the ^15^N_2_ bubble was completely dissolved. The % atom was directly measured in each microcosm bottle using a bench-top membrane-introduction mass spectrometry (MIMS, HPR-40 DSA). We filtered the bottle content through pre-combusted (450 °C, 4.5 h) GF/F Whatman filters, following incubation for 24 in running seawater to maintain the ambient temperature. The filters were dried at 60 °C overnight and analyzed using a CE Instruments NC2500 elemental analyzer interfaced with a Thermo-Finningan Delta Plus XP isotope ratio mass spectrometer (IRMS). The natural isotopic abundance of ^15^N_2_ (that is, seawater without the addition of ^15^N_2_) was measured and subtracted from the corresponding samples. A standard curve of acetanilide (C_8_H_9_NO) was generated before the measurements to determine the nitrogen mass of the samples (R^2^ > 0.99). The detection limit for ^15^N_2_ fixation was 0.02 nmol N L^-1^ d^-1^.

## Results and discussion

### Heterogenous microbiota is associated with tar patties

Using both amplicon sequencing of the 16S rRNA gene and metagenomics, we uncovered a large taxonomic diversity of bacteria and archaea that were associated with tar. Despite the considerable heterogeneity of prokaryotic communities associated with tar, they generally clustered based on the collection date (**Supplementary Figures S2 and S3**). We note that this pattern does not necessarily follow the dynamics of the oil weathering or biofilm succession, but rather likely exemplifies the heterogeneity of tar patties that were subjected to different environmental factors. For some tar particles, we observed a marked separation between the microbiomes of the inner, solidified tar and the outer ones that at least partially represent the exterior biofilms (e.g., samples 3-6, **Supplementary Figures S2 and S3**). In the samples that were collected afterward, we focused on the exterior biofilms.

The tar-associated communities differed markedly from those of ambient water, degraded plant material and biofilms on plastics **(Supplementary Figures S2 and S3)**. Using the amplicon sequencing of the 16S rRNA genes, we identified 75 prokaryote orders and 146 genera whose abundance was significantly enriched in tar compared to the other samples (adjusted p-value < 0.05, **Figure 2, Supplementary Figure S4, Supplementary Tables ST1 and ST2**). At the order level, we found marked enrichment of numerous putative degraders of petroleum, including the anaerobic sulfate reducers such as Petrotogales and Desulfobacterales, as well as the chemotrophic Acidithiobacillales, Halothiobacillales, and Ectothiorhodospirales, among others. Key enriched genera included the OHCB such as *Alcanivorax, Marinobacter* and *Ketobacter*, and other hydrocarbon degraders including *Mycobacterium* (Liew et al., 2014) and various sulfate-reducing anaerobes (e.g., *Desulfatiglans,* SEEP-SRB1), and tolerant-non OHCB organisms such as *Marinobacterium* (Dellagnezze et al., 2018). Most interestingly, cyanobacteria were highly enriched in tar (3.8 log2-fold change of Cyanobacteriales, with prominent genera *Coleofasciculus* and *Phormidium*, **Figure 2**). We thus hypothesize that they are associated with biofilms on the exterior of these particles. Indeed, we often observed green-brownish overgrowth on the surface of tar patties, providing evidence for the occurrence of photosynthetic organisms (**Supplementary Figure S5**).

**Figure 2:**
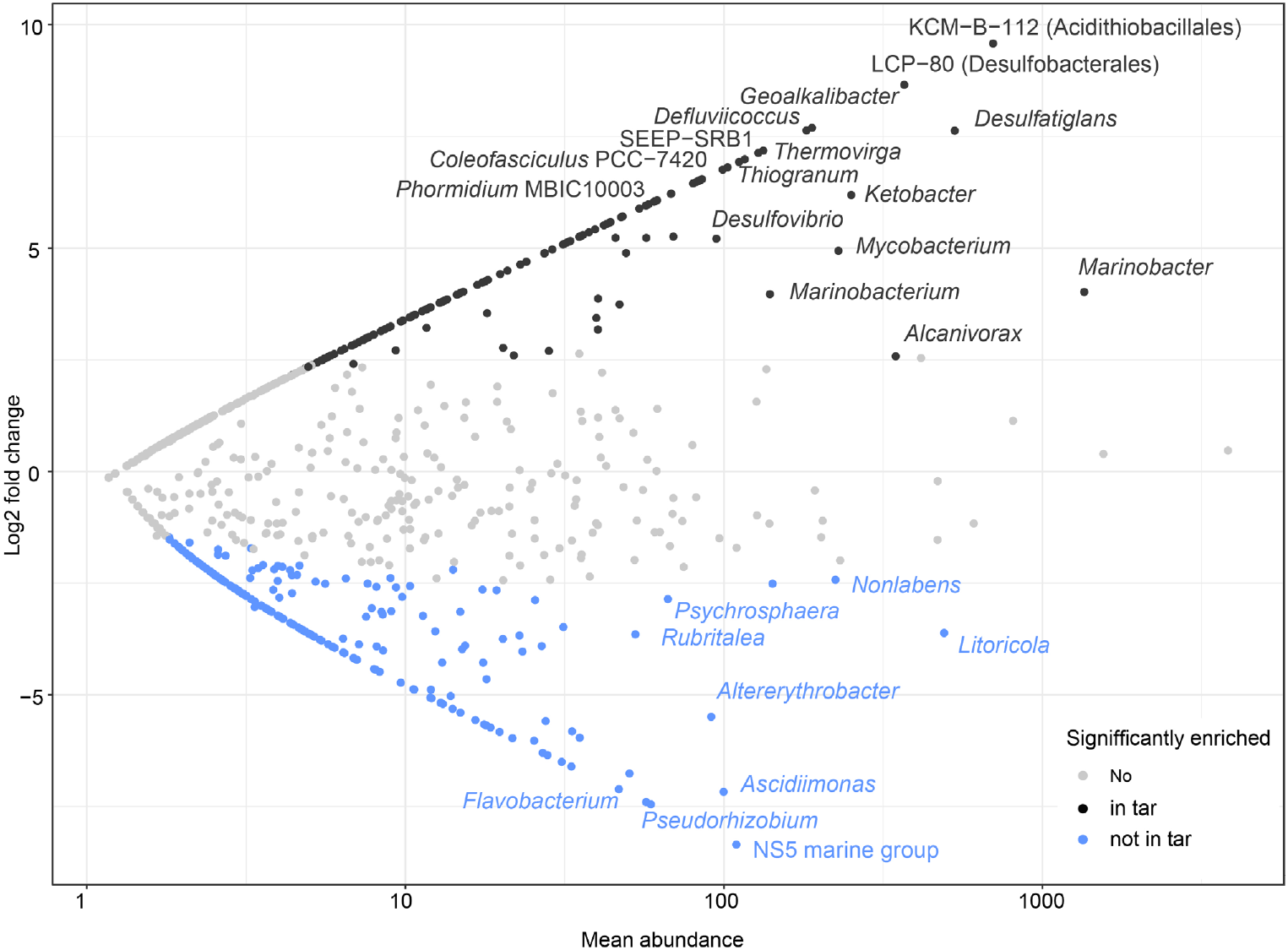
Enrichment of taxa in tar samples (black, positive log2 fold change), compared to biofilm, plastic and organic matter controls (genus level, DESeq2, significant when adjusted p-value <0.05). Only the most abundant differentially abundant genera are shown, the full list is shown in **Supplementary Table ST1**.

The deep shotgun sequencing of eight selected tar samples resulted in the curation of 79 medium to high-quality MAGs (**Supplementary Table ST3)**. Additional 5 MAGs were specific to plant material, representing the populations of *Actinoplanes brasiliensis*, *Brevundimonas* sp., *Pseudorhizobium pelagicum*, Pseudohongiellaceae CAILUG01 sp. and *Steroidobacter* sp., whereas no unique MAGs were associated only with plastic, where the dominant lineage was *Erythrobacter* sp. (78% read abundance). Hereafter we focus only on the tar-associated populations.

The diversity patterns based on metagenomics were similar to those resulting from the 16S rRNA gene amplicon sequencing, whereas some taxonomic inconsistencies were present due to the use of different reference databases (GTDB and Silva138) (**Figure 3**). For example, the prominent *Macondimonas* populations, identified by metagenomics, are referred to as Ectothiorhodospirales sp. by Silva138 taxonomy. Metagenomics highlighted the marked read abundance of mainly proteobacterial and mycobacterial obligate and facultative hydrocarbon-degraders, as well as large taxonomic diversity of anaerobic Desulfobacterota lineages, such as *Desulfoglaeba*, SEEP-SRB1c (C00003060) and Desulfatiglandaceae (**Figure 3**). Among the putative OHCB, the key clade detected by the metagenomics was UBA5335, to which the ubiquitous OHCB *Macondimonas* belong (Karthikeyan et al., 2019). Additional OHCB included *Marinomonas*, *Ketobacter* and *Mycobacterium*, among others. Following the results of the 16S rRNA gene amplicon sequencing, Cyanobacteria were among the prominent organisms associated with tar exterior, and comprised genera such as *Rivularia* and *Okeania* (**Figure 3)**. In samples collected on March 3^rd^ from the inner section of tar patties, we discovered a high read abundance of Syntropharchaeales, the eminent anaerobic oxidizers of short-chain alkanes (ANKA, 11.3-22.5% read abundance, **Supplementary Figure S6**) (Laso-Pérez et al., 2016; Wegener et al., 2022).

**Figure 3:**
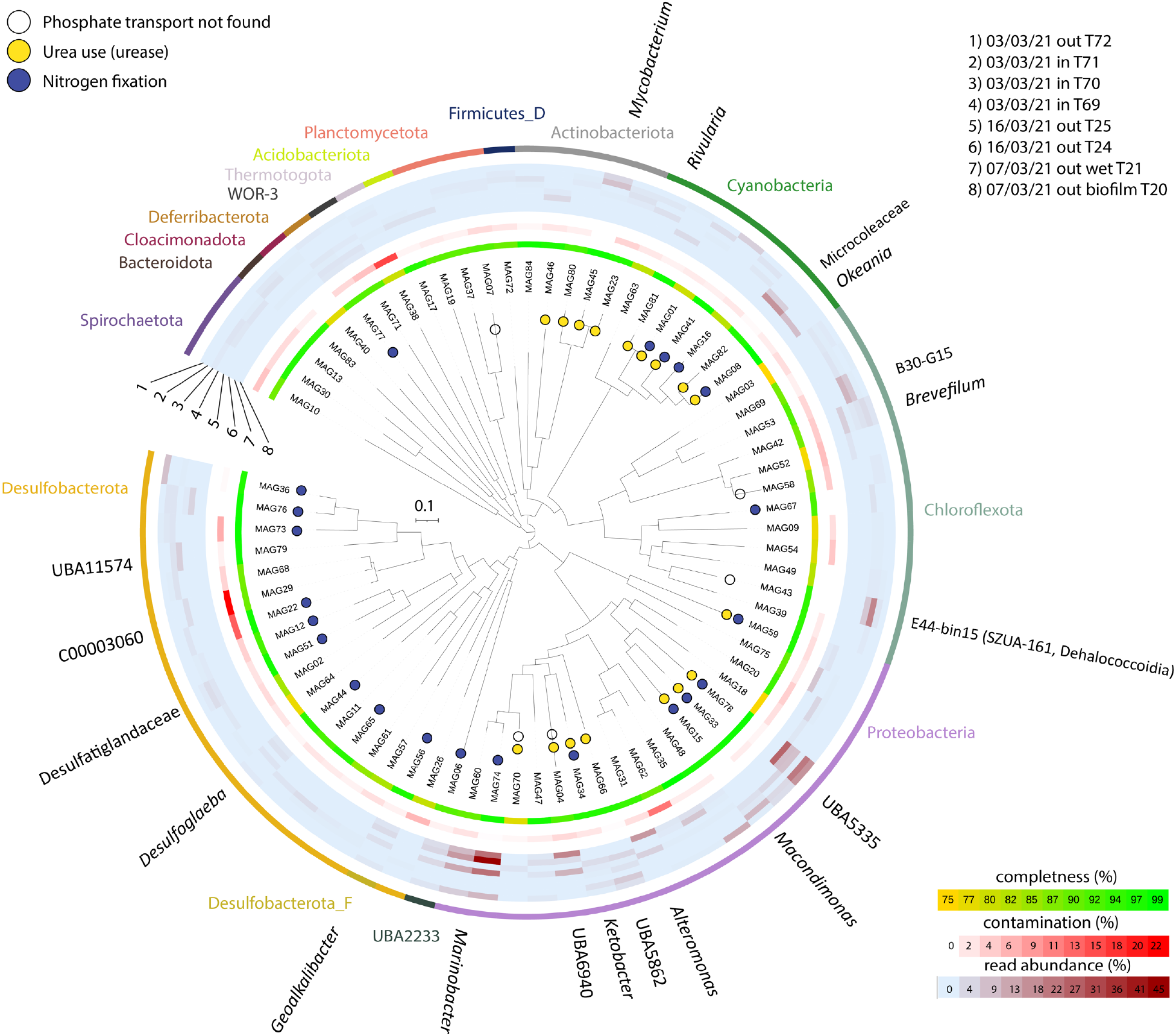
Taxonomy and abundance of metagenome-assembled genomes (MAGs) associated with seven tar patty samples from a nearshore pollution event in the southeastern Mediterranean Sea. Completeness, contamination (Checkm2) and read abundance statistics are shown. The FastTree tree is based on the alignment of GTDB marker genes, produced by GTDB-tk. Genus-level taxonomy for key lineages is shown. Only bacteria are included in this tree, for archaeal diversity and phylogeny see **Supplementary Figure S6**.

### Tar patties are hotspots of synergistic carbon and nutrient cycling

Metagenomics suggests that the tar-associated bacteria and archaea could collectively catabolize a broad range of organic substrates, via oxidation using electron acceptors such as oxygen, sulfate, or nitrate, but also ferment various organics (**Supplementary Figures S7 and S8**). No MAGs encoded all the denitrification steps, indicating that full denitrification can be performed only conjointly (**Supplementary Figure S8**). The sulfur reduction appears to be most prominent in the potentially anoxic tar interior, where most Desulfobacterota reside (**Figure 3, Supplementary Figures S7 and S8**). These organisms can couple sulfate reduction to oxidation of various petroleum-derived organic substrates (see below), as well as small organics such as acetate (Davidova et al., 2018). Some prominent anaerobic taxa, such as Dehalococcoidia (e.g, MAG43), appear to be acetogens, encoding the carbon monoxide dehydrogenase/acetyl-CoA synthase, as well as the genes needed to complete the tetrahydrofolate branch of the Wood-Ljungdahl pathway (Adam et al., 2018), but also ferment organics such as sugars, amino and fatty acids, which, in turn, can be derived from other community members. This example suggests that intricate metabolic interactions take place in the tar particles, as suggested previously for oil (Head et al., 2006).

In the ultraoligotrophic SEMS, key nutrients such as phosphorous and nitrogen are usually scarce. All but six MAGs encoded the high-affinity Pts phosphate uptake system, hinting a the competition for this resource. Uptake of inorganic nitrogen as nitrate appears to be limited, as only 15 MAGs encoded the assimilatory nitrate reductase (KEGG orthology: K10534), but AmtB ammonium transporters were widespread (found in 67 MAGs). Several abundant bacteria, including UBA5335 (*Macondimonas*) and Mycobacteriales, encoded the ABC urea transporters and the urease operon, as well as the guanidine degradation gene cluster (mainly in mycobacteria) (Schneider et al., 2020; Sinn et al., 2021) (**Figure 3**), in agreement with the hypothesis that the dissolved organic nitrogen could be a catalyst of hydrocarbon degradation (Ron and Rosenberg, 2014).

### Tar degraders may alleviate N limitation by fixing N_2_

Our assessment of MAG-encoded functions revealed that dinitrogen fixation is widespread among the tar-associated prokaryotes, as 26 out of 84 MAGs encoded the nitrogenase gene cluster (**Figure 3**). Diazotrophs were found across the clades of most prominent prokaryotes in our samples, e.g., *Marinobacter* (up to 45% read abundance in sample 14), UBA5335 (*Macondimonas*, up to 26% read abundance in sample 9), various Desulfobacterota (*Desulfosudis*, *Geoalkalibacter*, *Desulfoglaeba alkanexedens,* etc.), Cyanobacteria (*Okeania*, *Lyngbia*, Geitlerinemaceae genus LEGE-11467 and Spirulinaceae FACHB-1406), and the poorly-studied lineages such as UBA2233. All the Syntropharchaeales were identified as diazotrophs. These findings are in line with previous evidence of widespread potential to fix N_2_ among hydrocarbon degraders (Dashti et al., 2015; Foght, 2010), in particular, in the ubiquitous *Macondimonas* (Karthikeyan et al., 2019). Moreover, we highlight the potential for diazotrophy in anaerobic tar hydrocarbon degraders, in line with the widespread potential to fix N_2_ in natural deep-sea hydrocarbon seeps (Dekas et al., 2018, 2009; Dong et al., 2022; Metcalfe et al., 2021). Rate measurements in incubations suggest that tar amendments enhanced the N_2_ fixation rates in the SEMS coastal water (0.050±0.009 nmol N L^-1^ d^-1^ following tar addition, as opposed to 0.030±0.004 nmol N L^-1^ d^-1^ in controls, t-test, p<0.05), providing an additional line of evidence for stimulation of diazotrophy by oil-derived hydrocarbons. This ability to fix N_2_ may alleviate the marked limitation of hydrocarbon usage by nitrogen (Ron and Rosenberg, 2014; Rosenberg et al., 1998), in particular, in the ultraoligotrophic SEMS.

### Most tar-associated prokaryotes can degrade petroleum hydrocarbons

Our results suggest that tar patties are hotspots of hydrocarbon degradation, as various microbes encoded functions needed to degrade oil hydrocarbons, including (poly)aromatics and alkanes, both aerobically and anaerobically (**Figure 4, Supplementary Figure S9**). We detected the presence of these hydrocarbons, primarily a large range of n-alkanes from C_5_ to C_42_, aromatics and polyaromatics, using gas chromatography-mass spectrometry (GC-MS) analyses of oil that reached the shore on day one of the pollution event (**Supplementary Note 1**). We found the aerobic long-chain alkane monooxygenases and hydrolases mainly in proteobacterial Pseudomonadales (e.g., *Marinobacter*, *Marinobacterium*, *Ketobacter*), Enterobacterales (*Alteromonas*, *Pseudoaslteromonas*) and Mycobacteriales (*Mycobacterium*, *Gordonia*). Anaerobic degradation of alkanes via 1-methylalkyl (alkyl) succinate synthase (AssA) and alkane C2 methylene hydroxylase (AhyA) (Shou et al., 2021; Stagars et al., 2016) was linked to Desulfobacterota (e.g., Desulfatiglandales, UBA8473, and Adiutricales lineages) and Chloroflexota (Anaerolineae B4-G1, Dehalococcoidia SZUA-161 and UBA2777 lineages), and more rarely to Syntropharchaeales, Planctomycetota SM23-33, and Beggiatoales (**Figure 4, Supplementary Figure S9**). The degradation of aromatics and polyaromatics was widespread among the abovementioned lineages, but also was prominent in Acidimicrobiales (Actinobacteriota), and, more interestingly, in most tar-associated cyanobacterial lineages (benzene carboxylase in Geitlerinemaceae J055, *Lyngbia*, *Okeania*, *Rivularia* and Spirulinaceae ESFC-1; naphthalene carboxylase in *Rivularia*, which also encoded AlkB; as well as the aerobic naphthalene 1,2-dioxygenase alpha in Geitlerinemaceae J055). These findings hint that tar-associated cyanobacteria are likely not only tolerant to hydrocarbon pollutants but also can play a direct role in their remediation, that is, phytoremediation (Gupta et al., 2019). This is in line with the previous findings showing the ability of some cyanobacteria to degrade petroleum hydrocarbons (Raghukumar et al., 2001). Indeed, *Rivularia*, which appears to have the most remediation capacities, can be prominent in oil-polluted hydrosphere (Abed et al., 2014), and even in plastisphere (Zettler et al., 2013).

**Figure 4:**
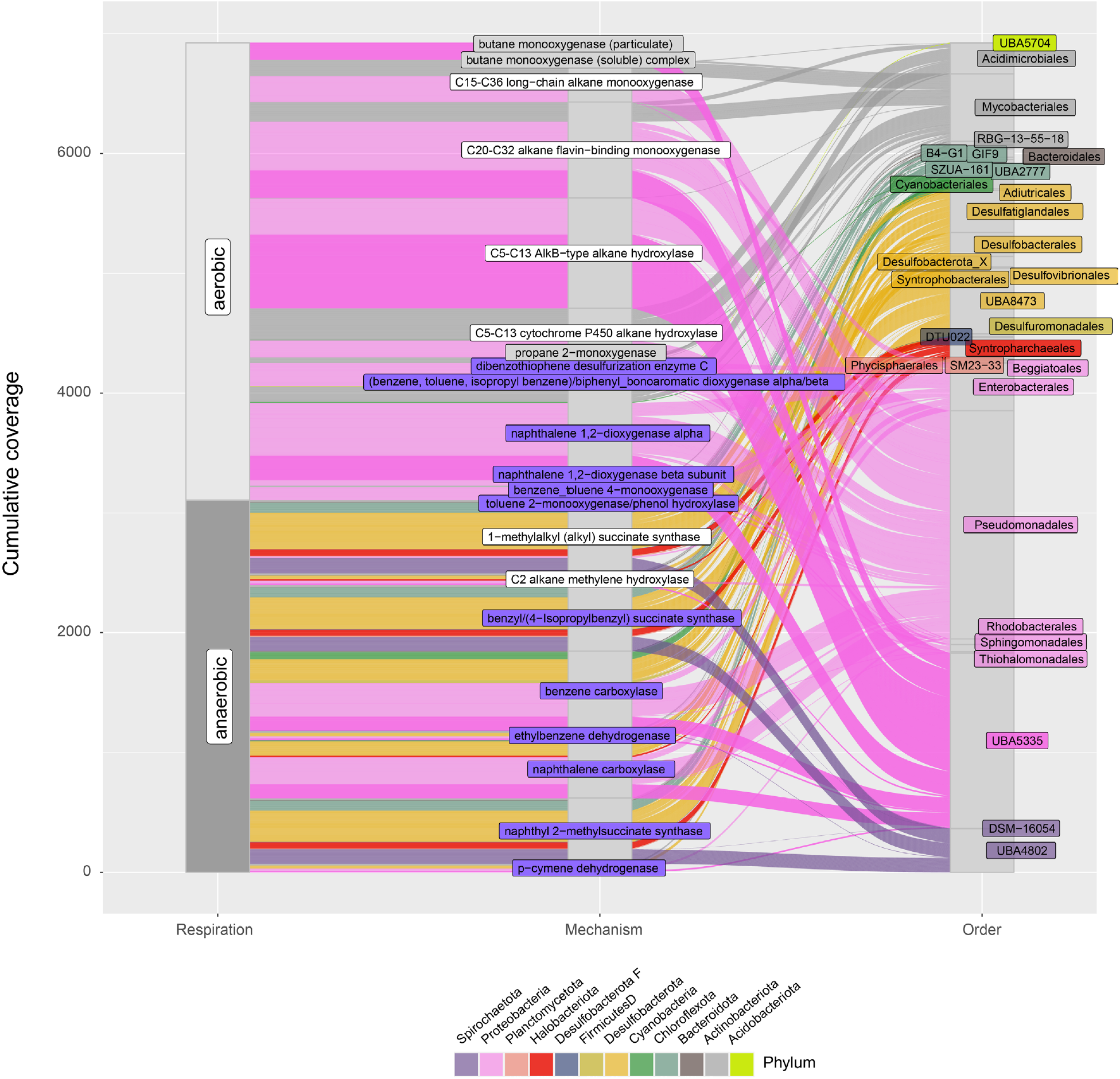
Relative abundance and taxonomy of key enzymes involved in the degradation of tar hydrocarbons, based on the CANT-HYD classification. Short-chain alkanes (grey), medium to long-chain alkanes (grey) and aromatics (purple) degradation pathways are highlighted, and both aerobic (e.g., Proteobacteria, Actinobacteria and Mycobacteriales) and anaerobic (e.g., Deuslfobacterota and Spirochaetota). The Syntropharchaeales acyl-CoM reductase for anaerobic butane oxidation is not shown because these AcrA proteins are not included in the CANT-HYD database.

### Short-chain alkanes may fuel tar-associated microbes

Whereas short-chain alkanes are likely a minor fraction of oil- and tar-derived hydrocarbons (**Supplementary Note 1**), our data suggest that these energy-rich compounds fuel some tar-associated microbes, potentially through cryptic cycling. Metagenomics supports this hypothesis, indicating that short-chain alkanes were potential carbon and/or energy sources for the dominant *Macondimonas*-like bacteria (clade UBA5335), mycobacteria *Mycobacterium* and *Gordonia*, as well as Syntropharchaeales ANKA (**Figure 4, Supplementary Figure S9 and Supplementary Table ST4**). CANT-HYD analyses identified the *bmoA* genes encoding the particulate butane monooxygenases in UBA5335 MAGs 15, 78 and 33; *bmoX* encoding the soluble butane monooxygenase, as well as *prmA* encoding the key subunit of the propane 2-monooxygenases in mycobacterial MAGs 45 and 80. UBA5335 encoded their particulate hydrocarbon monooxygenases as *phmCAB* (putative *bmoCAB*) operons, whereas additional copies of the *phmC* gene were present, suggesting a broader range of substrates affinities (Awala et al., 2021; Rubin-Blum et al., 2017; Wang et al., 2017) (**Supplementary Figure S10**). Yet, the UBA5335 PhmA/C sequences did not belong to the same clades as those of the known degraders of short-chain alkanes, indicating a distinct evolutionary path, and were highly aberrant to those of methane oxidizers, hypothetically indicating limited affinity to methane (**Supplementary Figures S10 and S11**).

Our data suggests that the abundant UBA5335 (*Macondimonas*) can use various short-chain alkanes as a carbon source and for energy conservation. Their MAGs encoded several short-chain-alkane-CoA synthases, as well as pathways to assimilate the activated products of C4, and potentially C2 and C3 alkane oxidation. This is supported by the presence of the putative butyryl-CoA dehydrogenase and enoyl-CoA hydratase, allowing further use of butane-derived carbon and energy (Rubin-Blum et al., 2017). Ethane and propane assimilation are also feasible, as UBA5335 encoded the glyoxylate shunt and methylcitrate cycle genes needed to use 2- and 3-carbon compounds (Ensign, 2006; Rubin-Blum et al., 2017). In turn, the methanol dehydrogenase and the key enzymes in serine and ribulose monophosphate (RuMP) pathways of formaldehyde assimilation were missing, indicating that methane is not used by these bacteria (missing in serine cycle: serine-glyoxylate aminotransferase, *sgaA*; hydroxypyruvate reductase, *hprA*; glycerate 2-kinase, *gckA*; missing in RuMP pathway: hexulose-6-phosphate synthase, *rmpA* and 6-phospho-3-hexuloisomerase, *rmpB*). This ability to use a broad range of petroleum hydrocarbons, including short- and long-chain alkanes, as well as (poly)aromatics, in a nitrogen-limited environment (see above the discussion of diazotrophy), likely underlines the success of this clade in the polluted oligotrophic hydrosphere.

In turn, *Gordonia* MAG80 and *Mycobacterium* MAG45, but not *Mycobacterium* MAG23, encoded the SmoXYB1C1Z soluble methane monooxygenase (sMMO)-like enzymes needed for using C2-C4 alkanes, likely propane and butane (Martin et al., 2014). The *Gordonia* SmoX was 100 % similar to that of propane-2-monooxygenase of *Gordonia mangrovi,* UniProt accession A0A6L7GMD1, whereas SmoX of *Mycobacterium* MAG45 (95.5% similarity to the *Mycobacterium dioxanotrophicus* SmoX UniProt accession A0A1Y0CHK1) belonged to an aberrant group of alkane-2-monooxygenases, also found in *Rhodococcus rubrer* (UniProt accession A0A098BII1) (**Supplementary Figure S12**). We identified the glyoxylate shunt genes in all the mycobacterial MAGs, but found the methylcitrate cycle genes only in *Gordonia*. The butyryl-CoA dehydrogenases were present (annotated as FadE25 by SEED): we identified them based on homology with previously-annotated mycobacterial butyryl-CoA dehydrogenases (Beites et al., 2021). All the tar-associated mycobacteria likely only partially rely on the short-chain alkanes, as these MAGs encoded a broad range of genes needed for catabolism of sugars, proteins, long-chain alkanes, mono- and poly-aromatics, as well as the degradation of more complex macromolecules, such as steroids and polystyrene (Cole et al., 1998; Johnston et al., 2010; Liu et al., 2023; Miner et al., 2009). The largest diversity of FadE proteins (EC 1.3.8.1) needed to degrade these complex macromolecules was in MAG23 (22), whereas MAGs 45 and 80 encoded 11. These mycobacteria can likely thrive under a broad range of redox conditions, capable, for example, of carbon monoxide and formate respiration using oxygen or nitrate as terminal electron acceptors, encoding the formate dehydrogenase O (EC 1.2.1.2 in all three MAGs), aerobic carbon monoxide dehydrogenase (EC 1.2.5.3, MAGs 23 and 80) and the respiratory nitrate reductase (EC 1.7.99.4, MAG45). Similar to *Mycobacterium smegmatis*, all these MAGs encoded the bidirectional NAD-reducing [NiFe] hydrogenase HoxFUYH (EC 1.12.1.2), potentially enabling tar-associated mycobacteria to produce dihydrogen under fermentative conditions (Berney et al., 2014). Thus, *Gordonia* and *Mycobacterium* appear to encode the traits needed to thrive both aerobically at the tar patty surface, and anaerobically in the interior.

In the interior of tar patties, the anaerobic degradation of short-chain alkanes was likely carried out by a distinct lineage of Ca. Synthroparachaeales. This archaea can fuel its metabolism with propane and butane, in partnership with a sulfate-reducing bacterium (Laso-Pérez et al., 2016; Wegener et al., 2022). For example, the Synthroparachaeales MAG25 encoded six copies of the alkane/methyl coenzyme M reductase alpha subunit AcrA protein, which clustered with those previously described in this clade (**Supplementary Figure S13**). We identified the genes needed for the short-chain alkane metabolism via β-oxidation pathway and the Wood-Ljungdahl pathway in most Synthroparachaeales MAGs, including the short-chain acyl-CoA dehydrogenase (*bcd*), enoyl-CoA hydratase (*crt*), 3-hydroxybutyryl-CoA dehydrogenase (*hbd*), acetyl-CoA acetyltransferase (*thl*), CO dehydrogenase/acetyl-CoA synthase (*cdhABC*), F420-dependent methylenetetrahydromethanopterin dehydrogenase (*mtd*, not in MAG25), 5,10-methenyltetrahydromethanopterin cyclohydrolase (*mch*), formylmethanofuran--tetrahydromethanopterin N-formyltransferase (*ftr*) and formylmethanofuran dehydrogenase (*fwdABC*) (Wang et al., 2021; Wegener et al., 2022). We did not find the *hmd* gene encoding 5,10-methenyl-H4MPT hydrogenase but found the *metFV* genes encoding the 5,10-methylenetetrahydrofolate reductase (EC 1.5.1.20), completing these pathways in the MAGs. The HdrABC complexes (*hdrABC* genes), needed to regenerate CoM-S-S-CoB were found, but we couldn’t trace most genes encoding the Fqo complex. We conclude that based on the high completeness of the abovementioned pathways in medium to high-quality MAGs, short-chain alkanes are the most likely substrates for the tar-associated Synthroparachaeales.

The partnership between these Synthroparachaeales and sulfate-reducing bacteria in these tar particles remains obscure. We did not find the canonical *Candidatus* Desulfofervidus auxilii syntrophs of Synthroparachaeales (Laso-Peréz et al., 2016), but observed a large diversity of Desulfobacterota, including the C00003060 SEEP-SRB1c, that can cooperate with the anaerobic methanotrophic (ANME) archaea (Skennerton et al., 2017). However, the FeGenie analyses identified extracellular electron transfer (EET) proteins only in other Desulfobacterota MAGs, mainly the Desulfovibrionaceae MAG26 (DFE_0461-4; DFE_0448-50) and *Geoalkalibacter* MAG56 (DFE_0461-4, OmcF, OmcS, porin-cytochrome (Pcc) protein complex) (Shi et al., 2014), which did not co-occur with Synthroparachaeales, indicating that they rather interact with inorganic particles or other prokaryotes. We detected a single candidate multiheme cytochrome for the extracellular electron transfer in the often-fragmented genomes of the tar-associated Synthroparachaeales, in MAG21, which contained 3 CHxxH motifs, as well as a non-cytoplasmic signal motif. Therefore, the partnership between Synthroparachaeales and sulfate reducers in petroleum pollution remains to be explored further.

### Micro- and macro-eukaryotes that colonize tar particles may promote nutrient cycling

Metagenomics-based interrogation of the small subunit ribosomal ribonucleic acids (SSU rRNAs) revealed that not only bacteria and archaea but various eukaryotes, including both protists and metazoans, were associated with tar and other debris (**Figure 5, see details at Figshare DOI:10.6084/m9.figshare.23498261**). Whereas plastics hosted mainly diatoms, following previous observations (Davidov et al., 2020), plant material was markedly enriched with fungi, in particular, *Corynespora* sp. (best hit *Corynespora cassiicola* UM 591, NCBI accession JAQF01001762.1, 232x coverage). Given that this fungal lineage is a ubiquitous plant pathogen (MacKenzie et al., 2018), its discovery in plant debris is not surprising.

**Figure 5:**
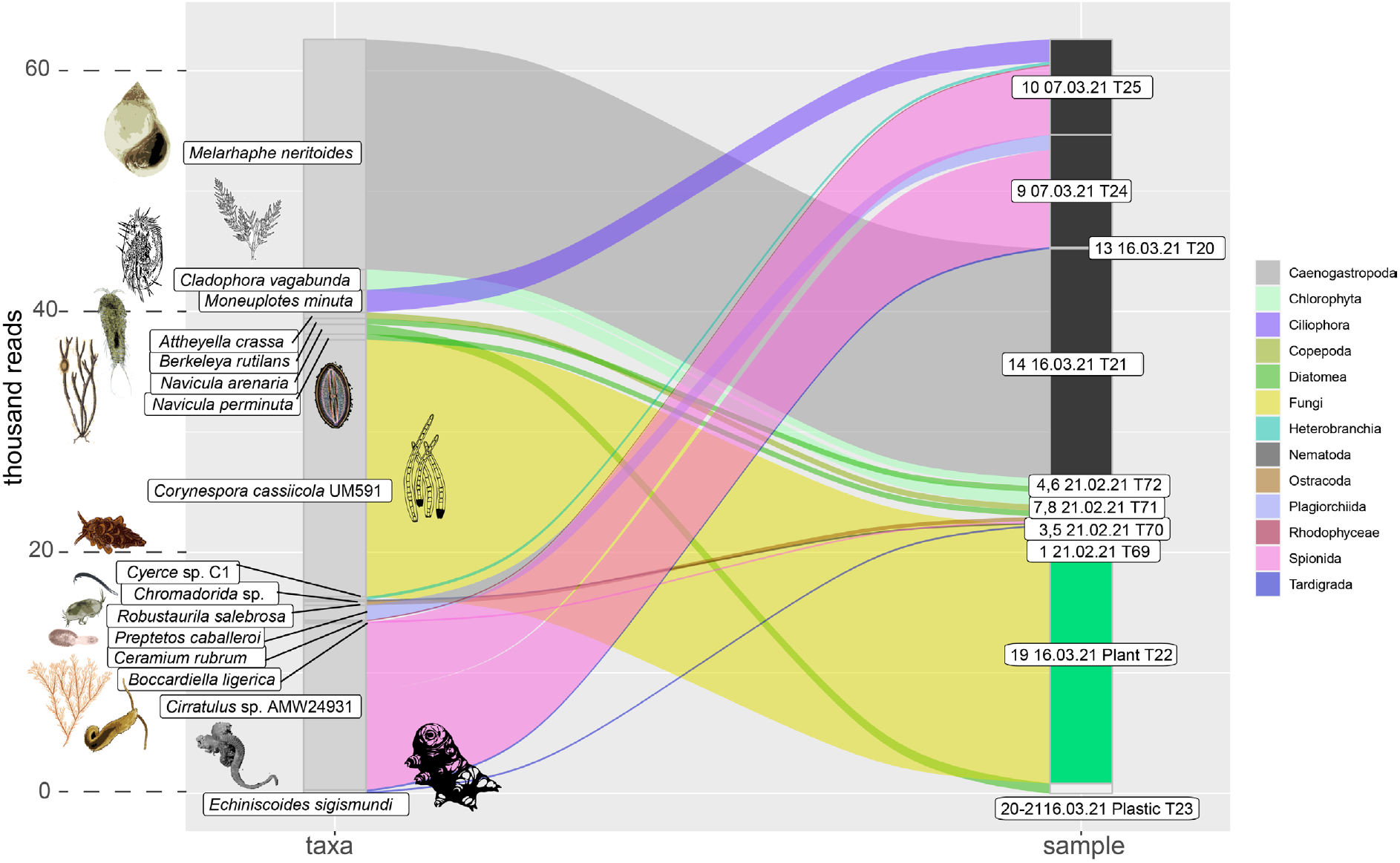
Taxonomy and read abundance of eukaryotes based on full-length 18S rRNA gene sequences that were assembled from reads that mapped to Silva 138 database, in tar (black), plant (green) and plastic (grey) samples. The paths are colored by high taxonomic level – we chose the most informative one for each lineage. The figure is illustrated with schematic, not up-to-scale, representations of taxa.

Tar patties carried a large diversity of associated eukaryotes (**Figure 5, Figshare DOI:10.6084/m9.figshare.23498261**). High-coverage eukaryote SSU sequences were mainly found in DNA from the fresh tar particles and likely represented potential living colonizers in the visible biofilms, but also could be derived from remnants that can randomly attach to the sticky surface (that is, environmental DNA). The abundant taxa included the common littoral fauna, such as *Cirratulus* annelids and *Melarhaphe* periwinkle gastropods, that were common in two samples collected on March 7th. Similar periwinkle species have been recently suggested as tar mat-associated heavy metals (Giraldes et al., 2022). The rarer taxa included *Moneuplotes* ciliates, green and red algae (*Cladophora* and *Ceramium*), diatoms, nematodes, sacoglossan sea slugs, as well as tardigrades (96.7% similarity to *Echiniscoides sigismundi* 18S rRNA gene sequence, NCBI accession GQ849021.1). For some taxa, such as foraminifera that were dominant in library T20 (specimen 13, tar biofilm), the SSUs did not assemble despite accounting for 62% of the reads mapped to the Silva database, and thus being highly abundant **(Figshare DOI:10.6084/m9.figshare.23498261**). This finding is supported by microscopic observations of numerous foraminifers attached to tar **(Supplementary Figure S5A**). We also observed the presence of *Lepas pectinata* on tar particles (different from those that were sequenced**, Supplementary Figure S5B,C**), confirming the previous observations of lepadid cirriped (goose barnacle) settlement on floating tar (Minchin, 1996).

While the interplay between the eukaryotes and their microbial neighbors remains to be elucidated, some interactions can be assumed, from grazing by bacteriovores to nutrient exchange. For example, the photosynthetic eukaryotes, such as diatoms, as well as red and green algae, may benefit from the accelerated nutrient turnover, in particular dinitrogen fixation, receiving new nitrogen from prokaryotic diazotrophs. In turn, algae may provide additional substrates for OHCB, releasing long-chain alkanes and other hydrocarbons into the phycosphere (Yakimov et al., 2022). Larger animals may recycle organic matter, and introduce new substrates for microbial metabolism, such as organic nitrogen. For example, crustaceans, as well as some gastropods, release urea (Conover et al., 1999; Mitamura and Saijo, 1980; Thibodeau et al., 2020), and the metabolic potential for urea use was widespread among the tar-associated prokaryotes, as described above. We assume that some of the prokaryotes described here may be associated with the larger animals, as gut- or epi-symbionts (Velasquez et al., 2023), providing an additional level of complexity to the biotic interactions in the tar communities.

## Conclusions

Our study suggests that tar patties are hotspots of microbial activity, both aerobic and anaerobic, which is largely fueled by oil-derived hydrocarbons, including short-chain alkanes. We hypothesize that as in large-scale spills (Valentine et al., 2010), priming of hydrocarbon-degrading communities in crude-oil pollution with energy-rich short-chain alkanes may occur also at smaller spill scales. While our analyses of the weathered oil did not target alkanes with carbon chains shorter than C5, we identified small quantities of C5 hydrocarbons, which could be catabolized by both anaerobic and aerobic prokaryotes. The cycling of short-chain alkanes is likely cryptic in the weathered particles, resulting in the undetectable release of these compounds into the hydrosphere. Yet, the rates of microbial short-chain alkane turnover and its contribution to the energy and carbon budgets of tar-associated communities remain to be determined.

In these tar-associated communities, the ability to use oil-derived hydrocarbons appears to be widespread not only in the OHCB, but also in other lineages, including Cyanobacteria, which can benefit from these energy-rich supplements, and/or detoxify the oil-derived molecules. Nitrogen fixation appears to give an advantage to key members of these communities, as nitrogen sources may be limited, especially in the oligotrophic eastern Mediterranean Sea. We also hypothesize that tar is a potential hotspot of mineral iron and phosphorous leaching, as the sticky tar patties that float on the sea surface can collect mineral-rich dust particles and given the EET in some taxa, e.g., Desulfobacterota, reduce extracellular minerals. Nutrient export from tar remains to be quantified.

We highlight the complexity of metabolic functions and handoffs in tar-associated communities, as well as in those that occupy other foreign particles in the marine environment. Tar weathering and subsequent colonization by biofilm-forming microbes, including photosynthetic bacteria, eukaryotic algae, protists and metazoans may increase the habitat complexity, and leverage the biotic interactions, yielding complex food webs. Similar to the phycosphere, metabolic handoffs in tar-associated communities likely involve the exchange of organic carbon, nitrogen and sulfur, sharing vitamins and cofactors, as well as the release and uptake of siderophores to capture trace metals (Smith et al., 2022). The tar-associated microbial populations may battle for these resources, given the occurrence of active competition mechanisms, such as the type VI secretion systems in the abundant *Marinobacter* MAGs 70 and 74 and common RTX toxin transport, found in 43 MAGs. These biotic interactions in tar, and petroleum-degrading communities in general, for example, symbioses between prokaryotes and eukaryotes, as well as the role of viruses, remain to be explored.

## Data availability statement

The raw metagenomic reads are available on NCBI as BioProject PRJNA981336.

## Author contributions

MR-B, TG-H and ER conceived the study and acquired funding. MRB and ER performed field sampling. YY and SM performed the molecular work. OR collected and documented the samples. MR-B performed bioinformatics. AA and IK performed GC-MS analyses of oil. MR-B wrote the paper with the contributions of all co-authors.

## Acknowledgments

This study is funded by the Israel Ministry of Energy (proposals 219-17-015 and 221-17-002) to MR-B and ER, the Israel Ministry of Science and Technology Grant (proposal # 001126) to MR-B and ER, the Israel Ministry of Environmental Protection (grant 195-5-1) to MR-B and ER and the National Monitoring Program of Israel’s Mediterranean Waters.

## Supplementary Material

### Supplementary Note 1: GC-MS analysis of the crude oil sample

The analyzed sample was collected in Bat-Yam Beach on 20/2/2021 (BY2). Water associated with the sample was dried by using magnesium sulfate. The dry sample was fractionated to saturates, aromatics, resins and asphaltenes by silica-gel column chromatography (SARA). The results are: Saturates =39.4%, Aromatics = 26.5%, Resins =3.6%, and Asphaltenes = 30.5% (Figure S14). Several grams of the sample were placed in a 40 ml vial sealed with septa and left at room temperature for 24 hours to allow the attainment of equilibrium between the liquid and gas phases. The headspace above the sample was then collected with a gas-tight syringe (100 μL) and injected into an Agilent 7890A GC coupled with an Agilent 5975C mass selective detector (GC-MS). The GC used a 30m·0.25mm·0.25μm DB-5MS fused silica column. Helium was used as a carrier gas at a constant rate of 1.5 ml/min. The temperature program was isothermal at 40°C for 14 minutes. The saturate and aromatic fractions were also analyzed by GC-MS. About 1 μL of each fraction was injected at split 10 and a constant injector temperature of 320°C. The temperature program used is 50°C for 5 minutes, a 5°C/min linear gradient up to 320°C with a hold at the final temperature (320°C) for 20 minutes. All other GC-MS parameters are the same as for the headspace analysis described above.

The total ion current (TIC) of the head space collected (Figure S15) demonstrated *n*-alkanes ranging from C_5_ to C_11_ with the presence of cycloalkanes. The TIC of the saturate fraction (Figure S16) demonstrated a large range of *n*-alkanes from C_10_ to C_42_ with the presence of iso- and cycloalkanes. Such observation indicated the analyzed hydrocarbon sample is likely a crude oil rather than tar or other refinery product. The very low content of *n*-C_10_ and the low abundance of relatively volatile low-molecular-weight aliphatic compounds in the range of *n*-C_10_ to *n*-C_16_ is indicative of evaporation and possibly some a certain degree of biodegradation that the sample has experienced. The loss of volatile compounds is usually coupled with a buildup of less volatile, high-molecular-weight, aliphatic compounds such as those observed in the range around *n*-C_34_. In addition, an increase in the UCM hump is visible which further supports the weathering experienced by the sample.

In the aromatic fraction (Figure S17), the very low content of naphthalene and C_1_-naphthalenes relative to longer, C_2_ and C_3_ naphthalene, is indicative of weathering, likely by water washing. Studies of oil spill weathering demonstrated the removal of all short benzenes takes place when oil evaporation reached ∼45% (Wang and Fingas, 1995). In the studied sample, the content of benzenes is very low which may further support weathering of the sample by evaporation.

---

**Figure S1:**
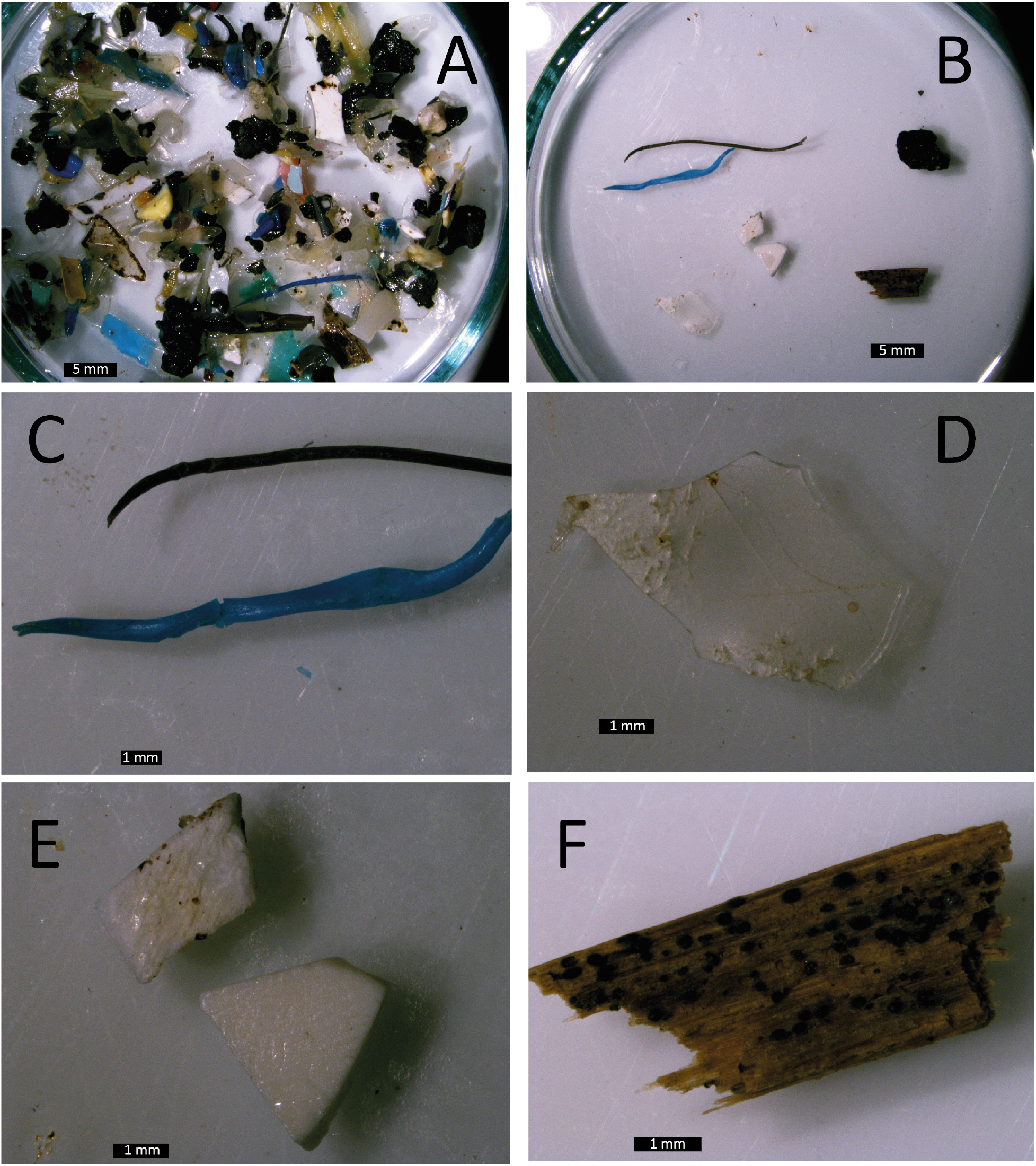
Debris collected from the tidal pool, following the tar pollution event (Tel Shikmona, Haifa, Israel, images taken on March 16, two weeks following the pollution event). Tar, plastic, and plant organics were collected (A) and separated for sequencing (B). The following were analyzed: plastic filaments (C); plastic sheets (D); microplastic fragments (squares, E); plant material (F).

**Figure S2:**
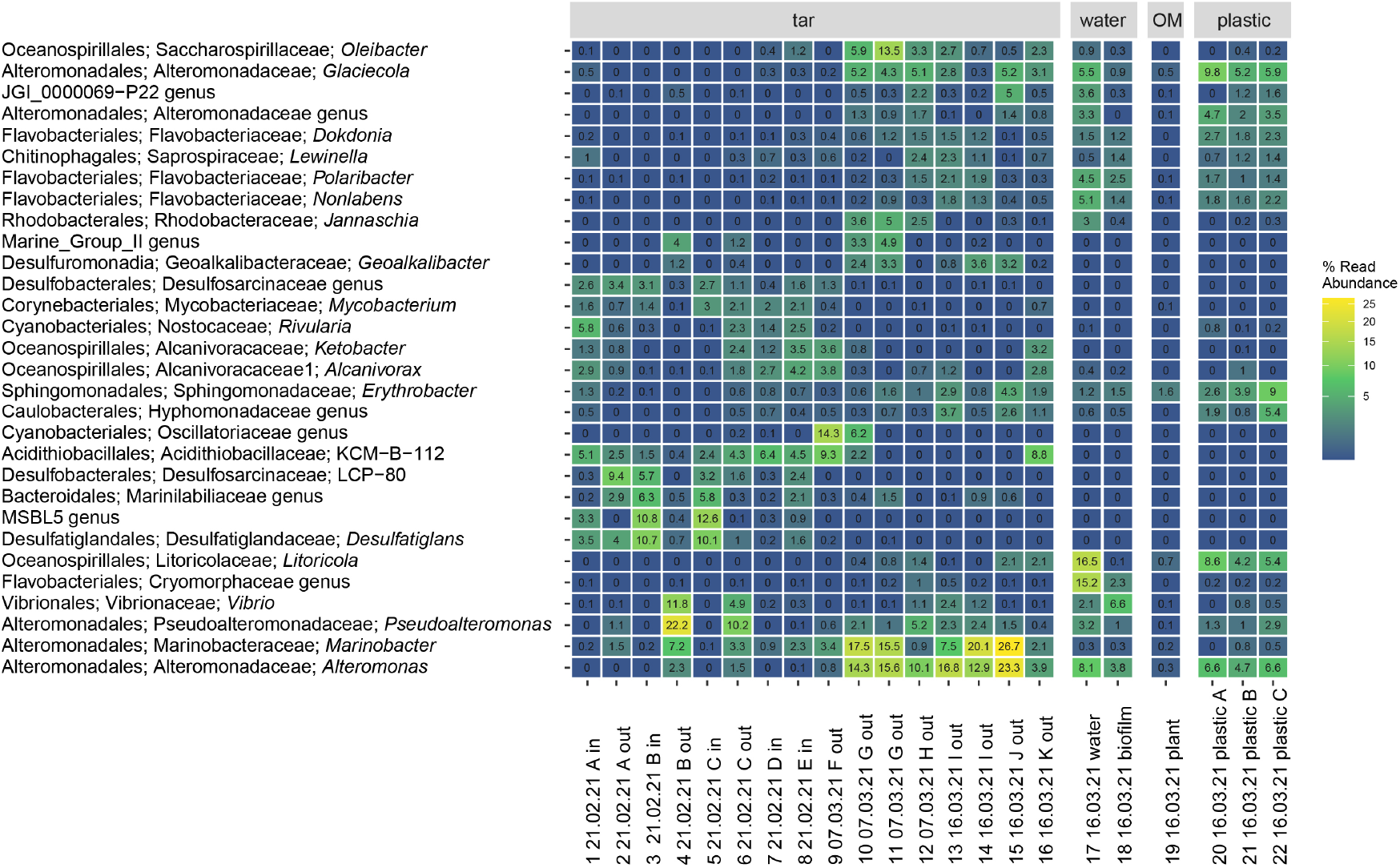
The relative read abundance of 30 key prokaryote genera (Silva v138 taxonomy) that were found in tar, ambient water and biofilm on its surface, plant-derived organics (OM) and microplastic samples of distinct morphologies - white squares (A), filaments (B) and sheets (C), based on the amplicon sequencing of the 16S rRNA gene. See Table 1 for sample details.

**Figure S3:**
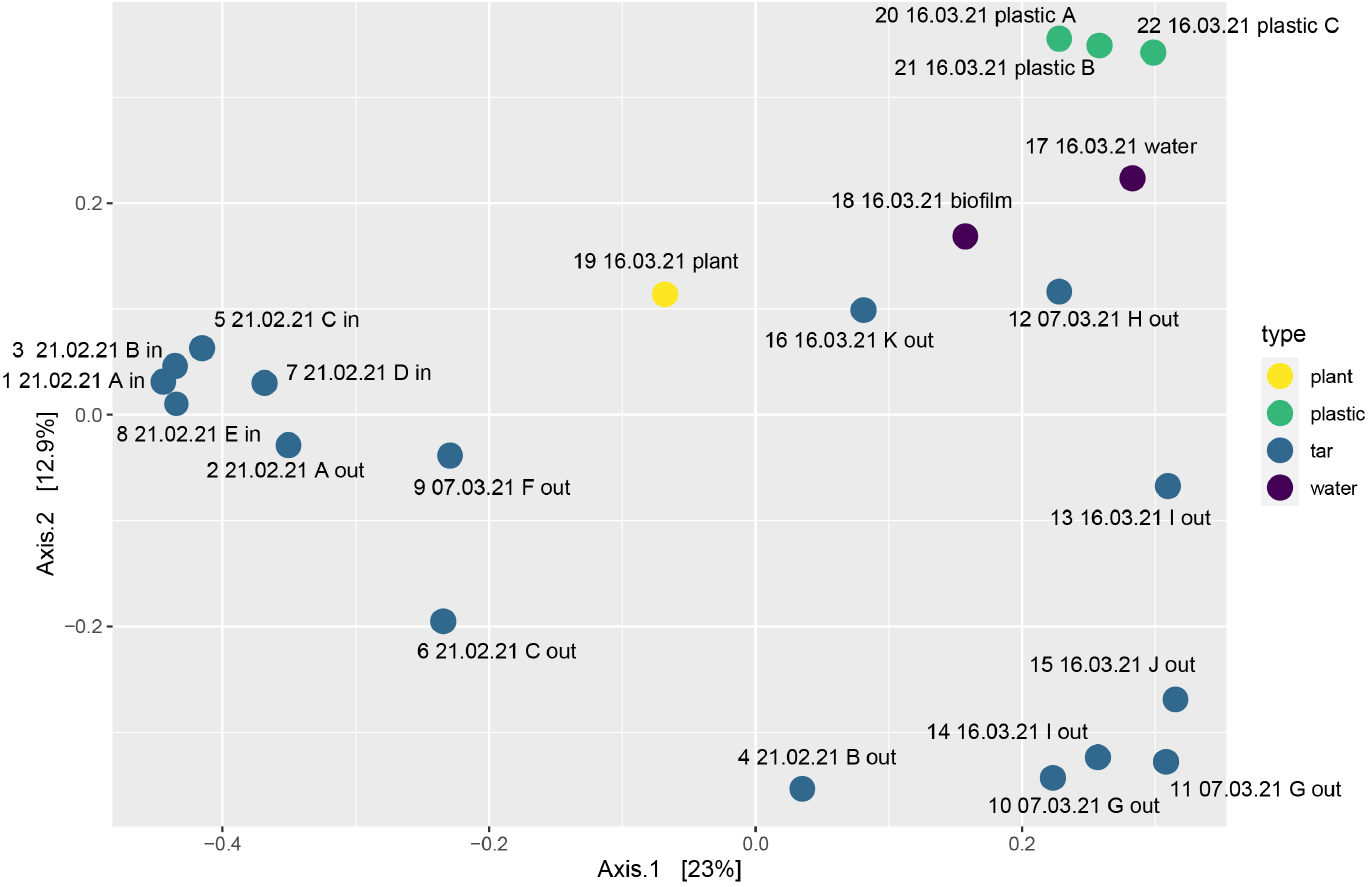
Principal coordinates analysis (PCoA, Bray-Curtis distance), showing the dissimilarities between tar, water, pant and plastic samples collected following the weathered oil pollution event in the southeastern Mediterranean Sea (February 2021). For samples description, please refer to Table 1 in the main text. The sample descriptors are as follows: running number, collection date, a letter indicating the whole tar sample, and in/out indicating the patty’s interior or exterior parts.

**Figure S4:**
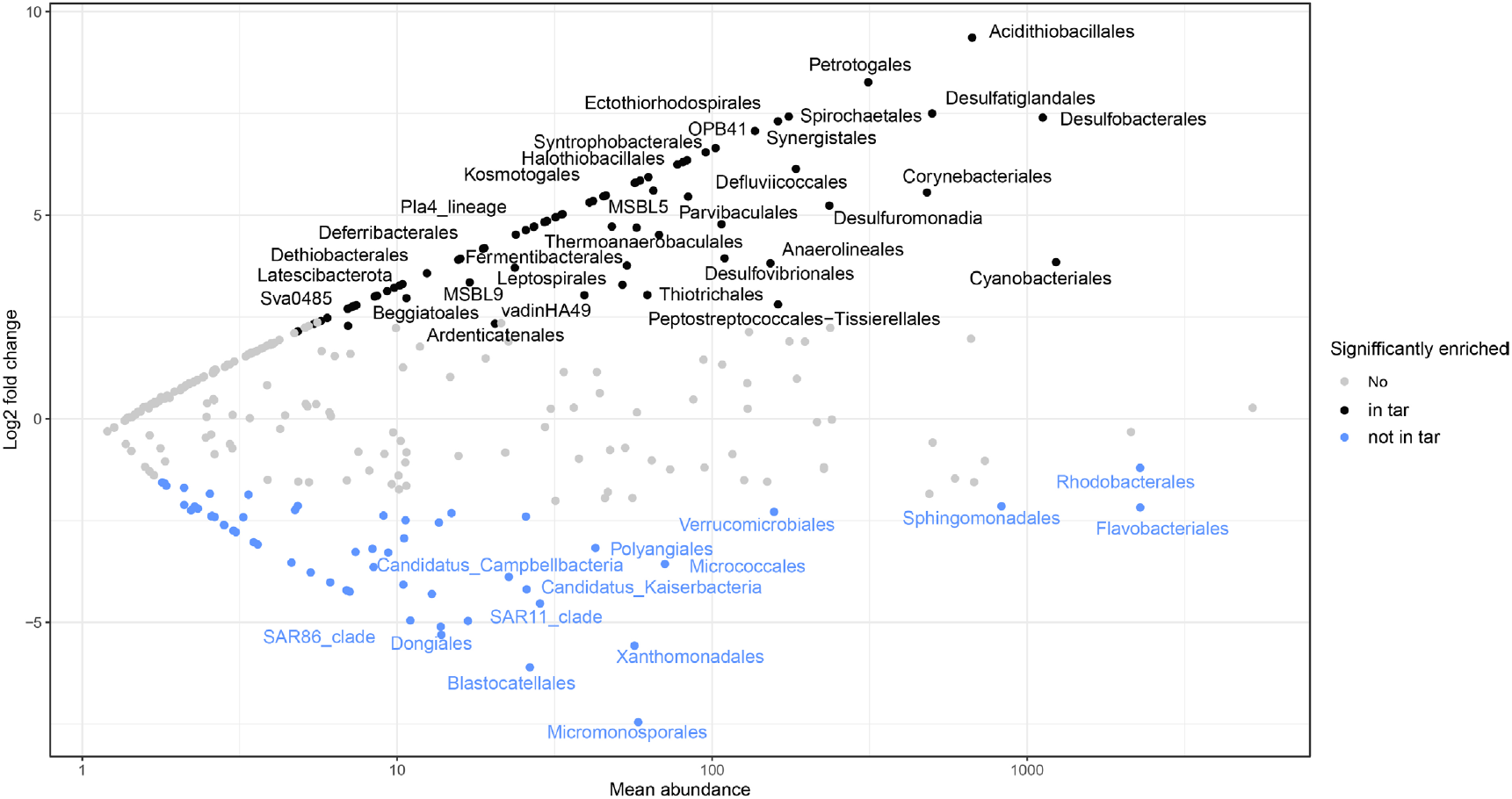
Enrichment of taxa in tar samples, compared to biofilm, plastic and organic matter controls (order level, DESeq2). Only the most abundant differentially abundant genera are shown, the full list is in **Supplementary Table ST2**.

**Figure S5:**
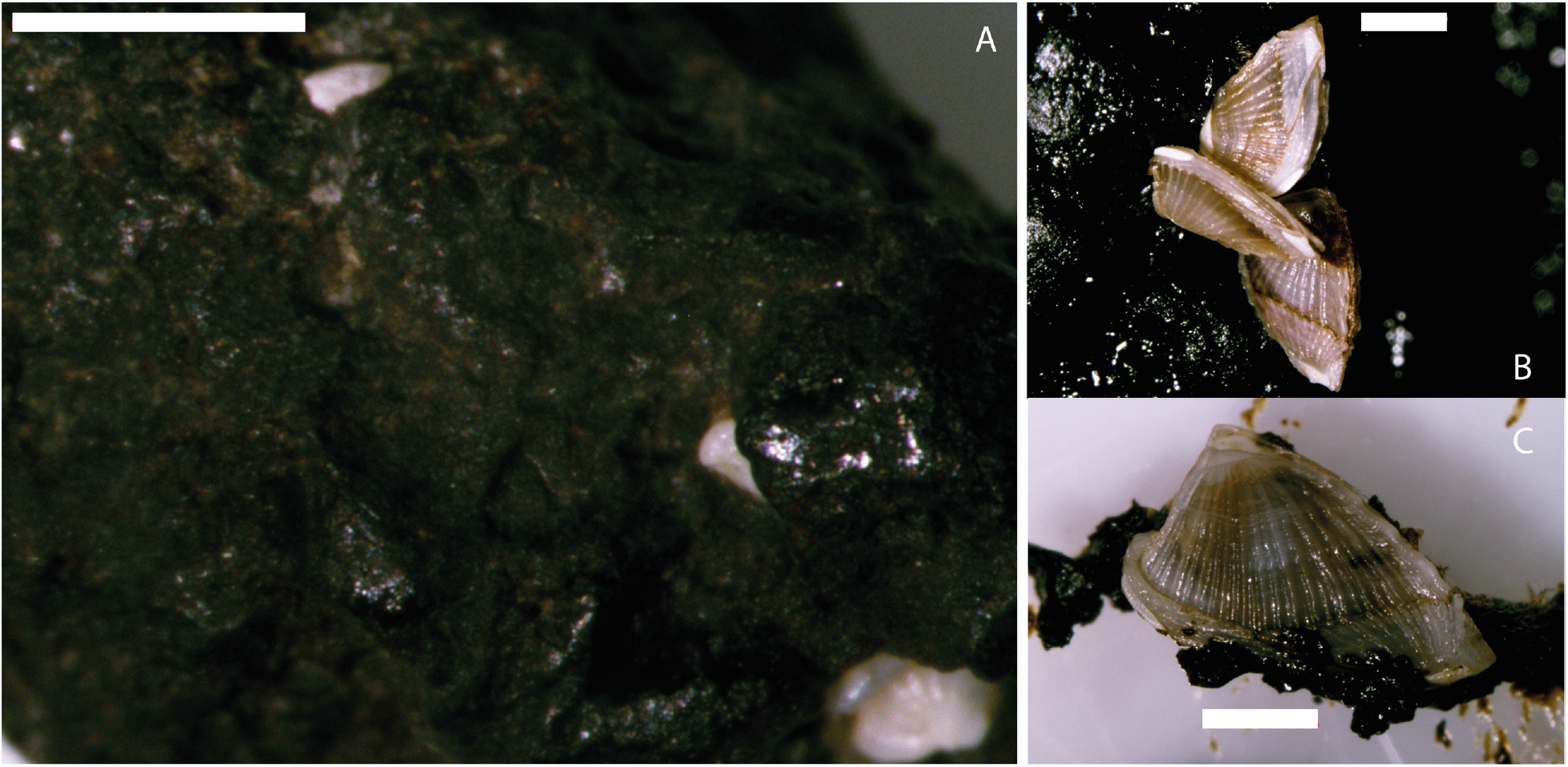
Metazoans and large protists colonizing tar surface. A) Foraminifers on a small tar particle collected on March 16^th^ 2021 from the tidal pool, alongside with tar that was sequenced. The particle is covered with a brownish biofilm. B) A cirriped *Lepas pectinata* colonizing tar particles collected in March 2021 (not included in the sequenced samples). C) *Lepas pectinata* on tar particles, green-brown overgrowth can be seen on the right side of panel B. Scale bar is 1 mm.

**Figure S6:**
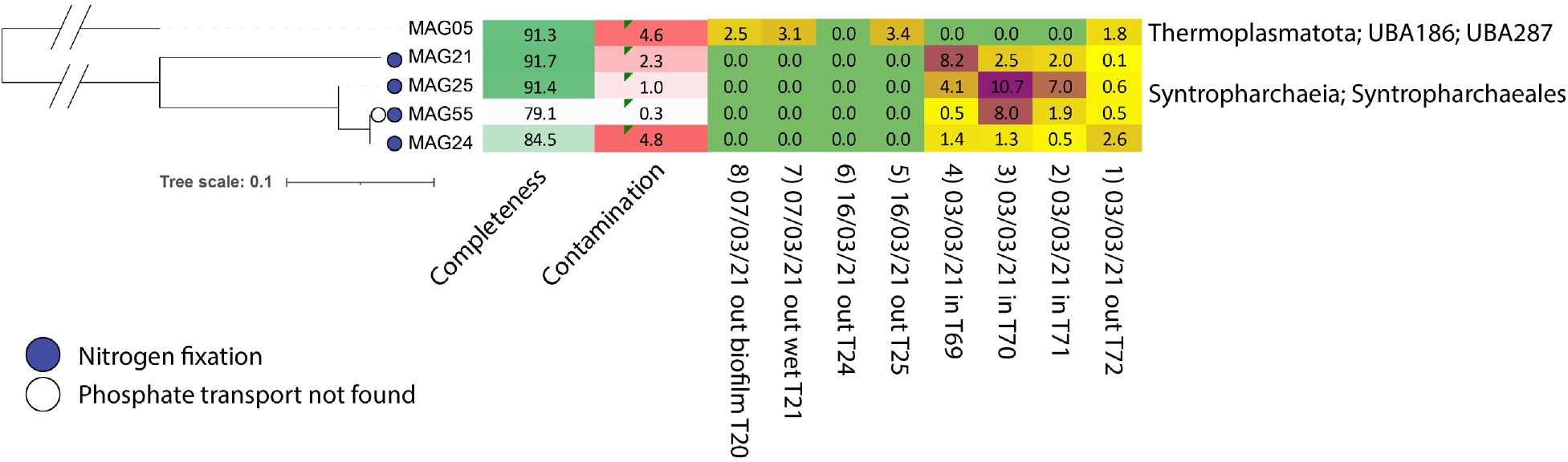
Phylogeny and abundance of archaeal metagenome-assembled genomes (MAGs) associated with seven tar patty samples from a nearshore pollution event in the eastern Mediterranean Sea. Completeness, contamination and read abundance statistics are shown. MAGs that encode the potential for N_2_ fixation are highlighted (red). The FastTree tree is based on the alignment of GTDB marker genes, produced by GTDB-tk.

**Figure S7:**
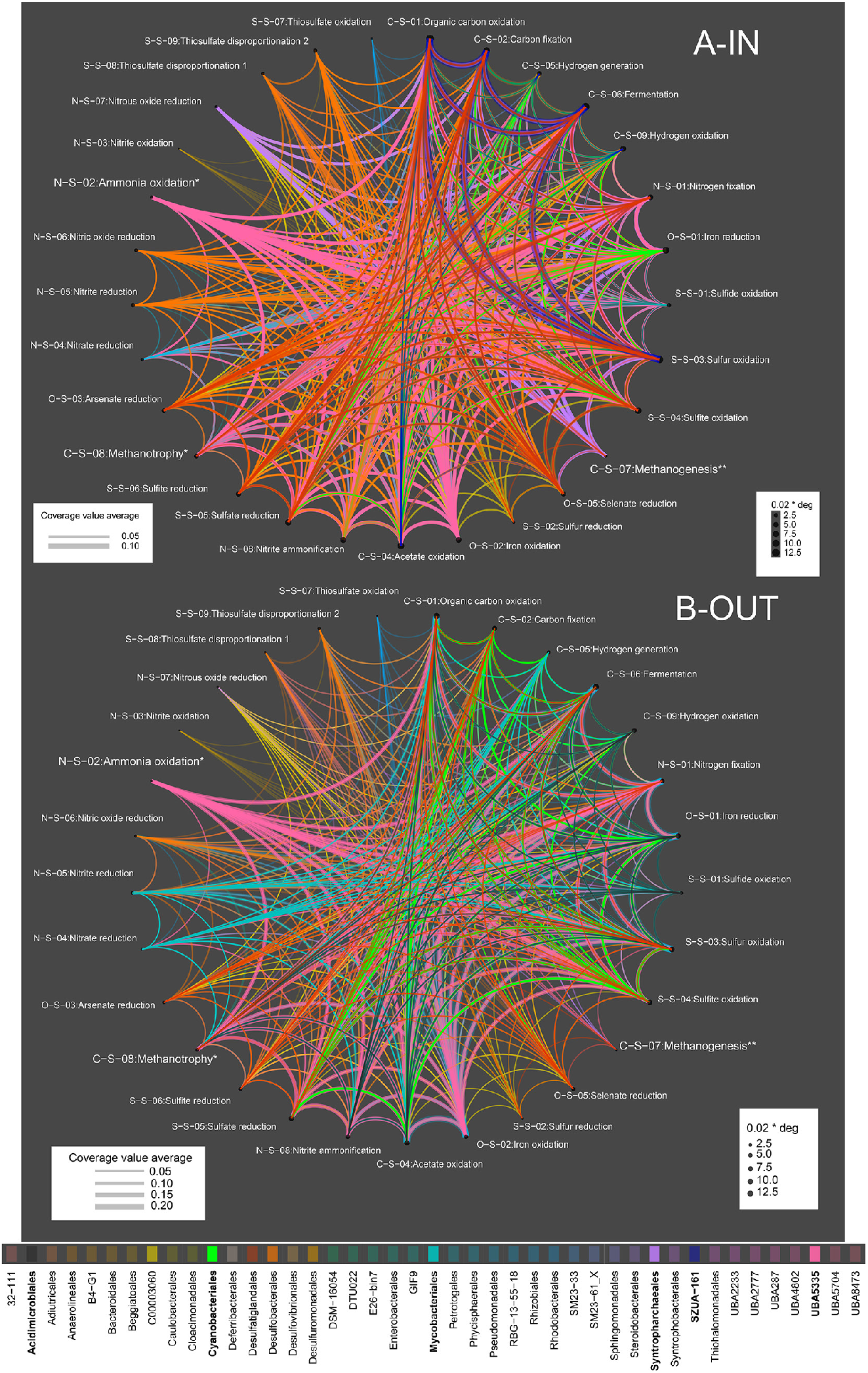
Metabolic functions and handoffs in tar-associated communities (A – inside, libraries T69-T71; B – outside, libraries T20, 21, 24, 25, 72), based on METABOLIC V4 reconstruction using the 0.9 pathway completeness threshold and order-level GTDB taxonomy. Path width is based on metagenomic coverage, vertex degree represents the number of connections (large vertices represent functions that displayed a large taxonomic diversity). * Methanotrophy and ammonia oxidation were identified based on the presence of ammonia/methane monooxygenases, yet our results indicate that these enzymes rather catalyze the oxidation of short-chain alkanes. ** Methanogenesis is identified by the presence of methyl-coenzyme M reductases (McrA-like proteins), yet our results indicate that these are rather alkyl-coenzyme M reductases AcrA, needed for the anaerobic oxidation of short-chain alkanes in Syntropharchaeales.

**Figure S8:**
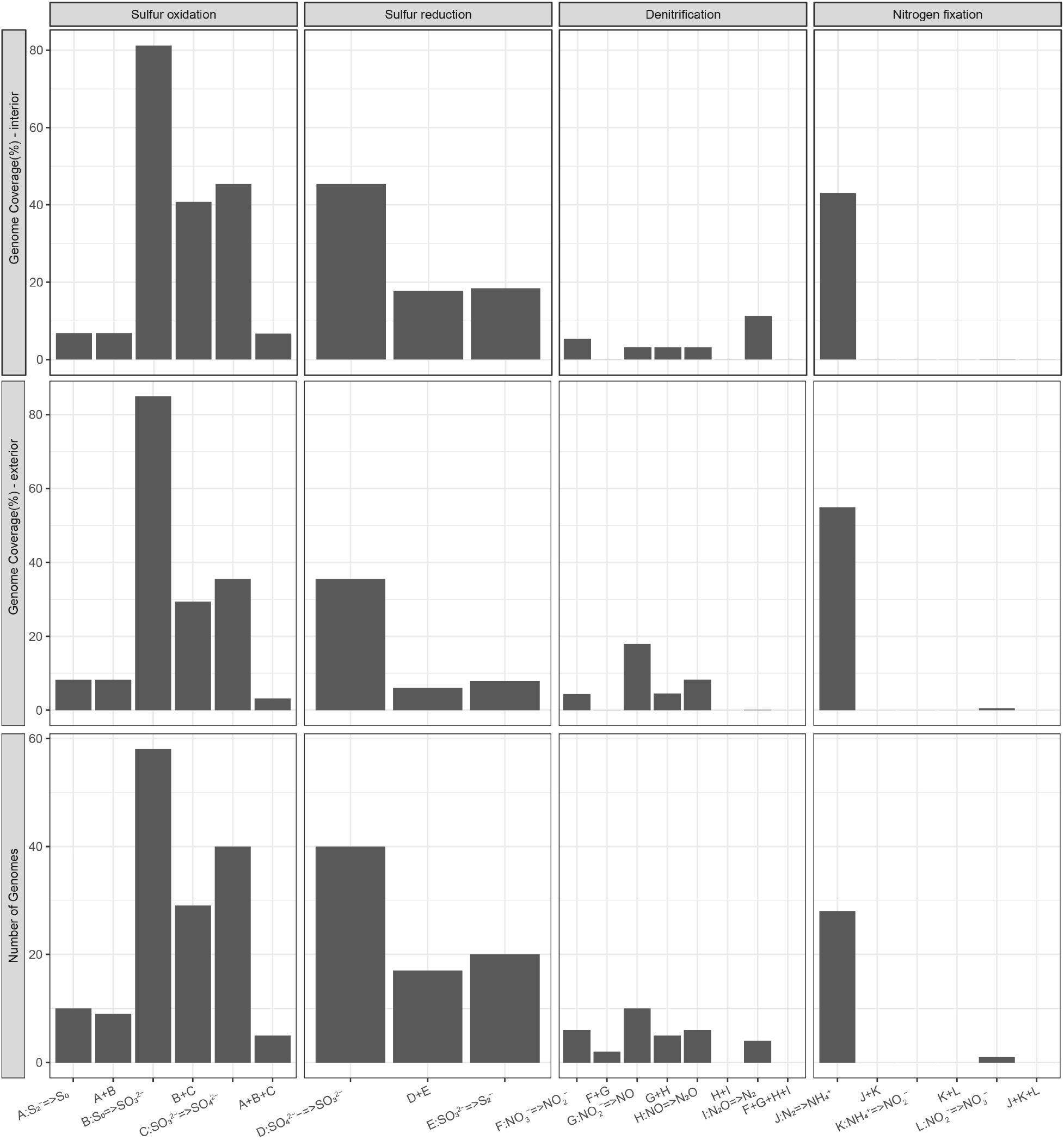
The sequential transformation of sulfur and nitrogen in microbial communities associated with tar interior and exterior, based on METABOLIC V4 reconstruction using the 0.9 pathway completeness threshold.

**Figure S9.**
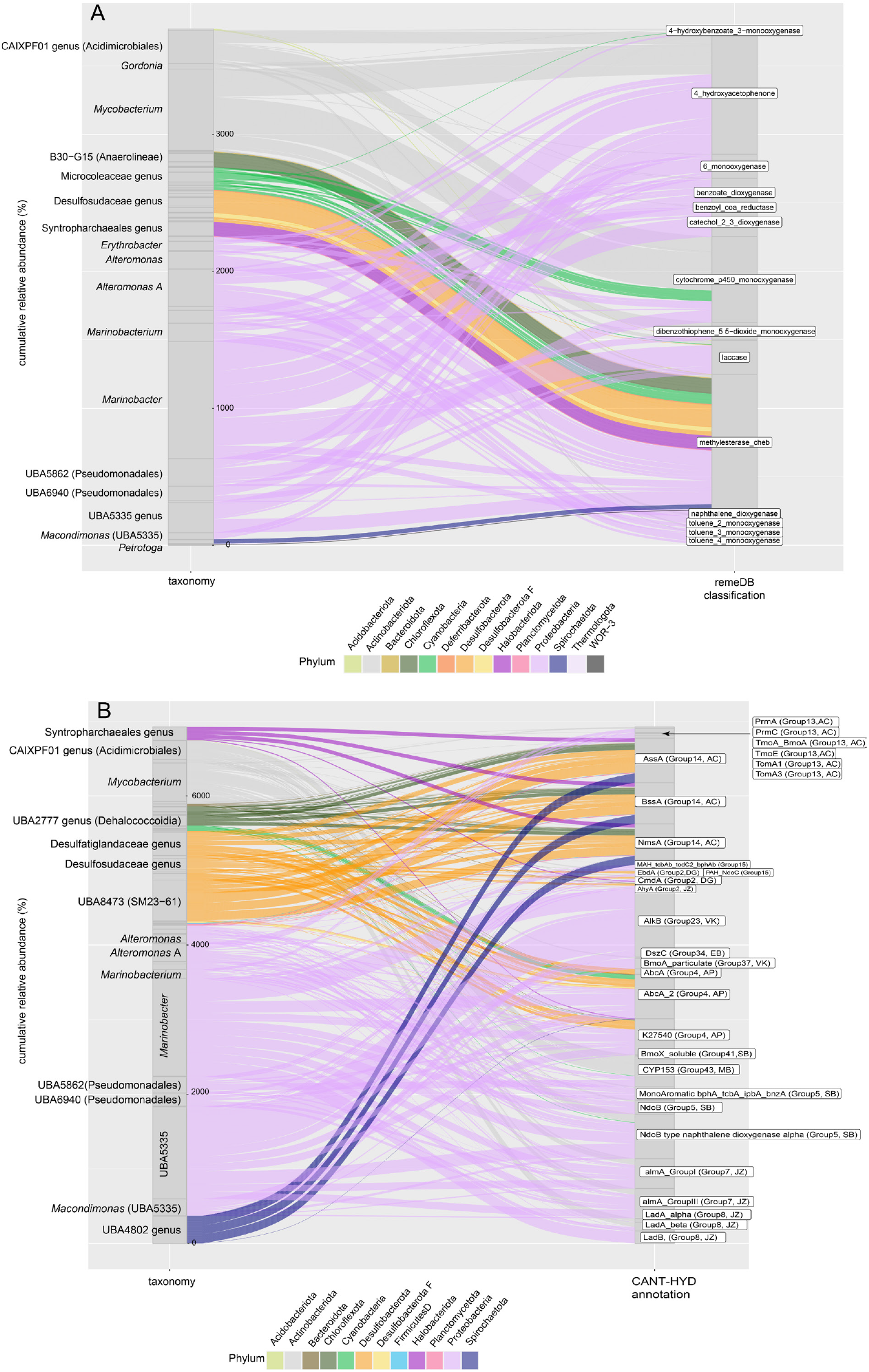
Relative abundance and genus-level taxonomy of key enzymes involved in the degradation of tar hydrocarbons and their owners, classified based on RemeDB (A) and CANT-HYD (B).

**Figure S10:**
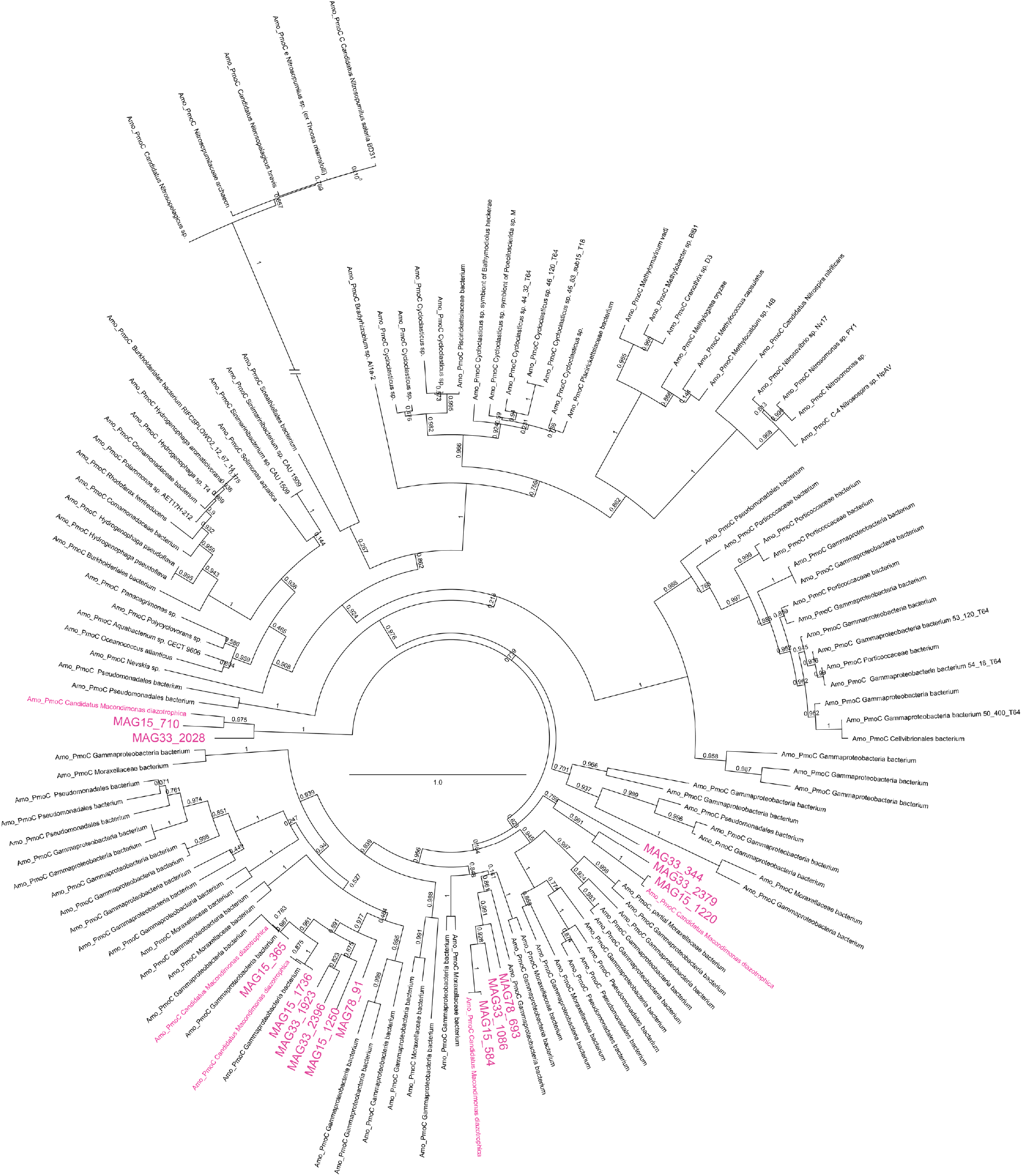
Phylogeny of PhmC/AmoC proteins in alkane-oxidizing UBA5335, including *Candidatus* Macondimonas, from tar pollution (purple), and related PhmC/PmmC/AmoC sequences from methane, short-chain alkane and ammonia oxidizers (FastTree). Branch bootstrap values are shown, and the scale represents the number of substitutions per site. The very long archaeal AmoC branch is truncated for ease of visualization.

**Figure S11:**
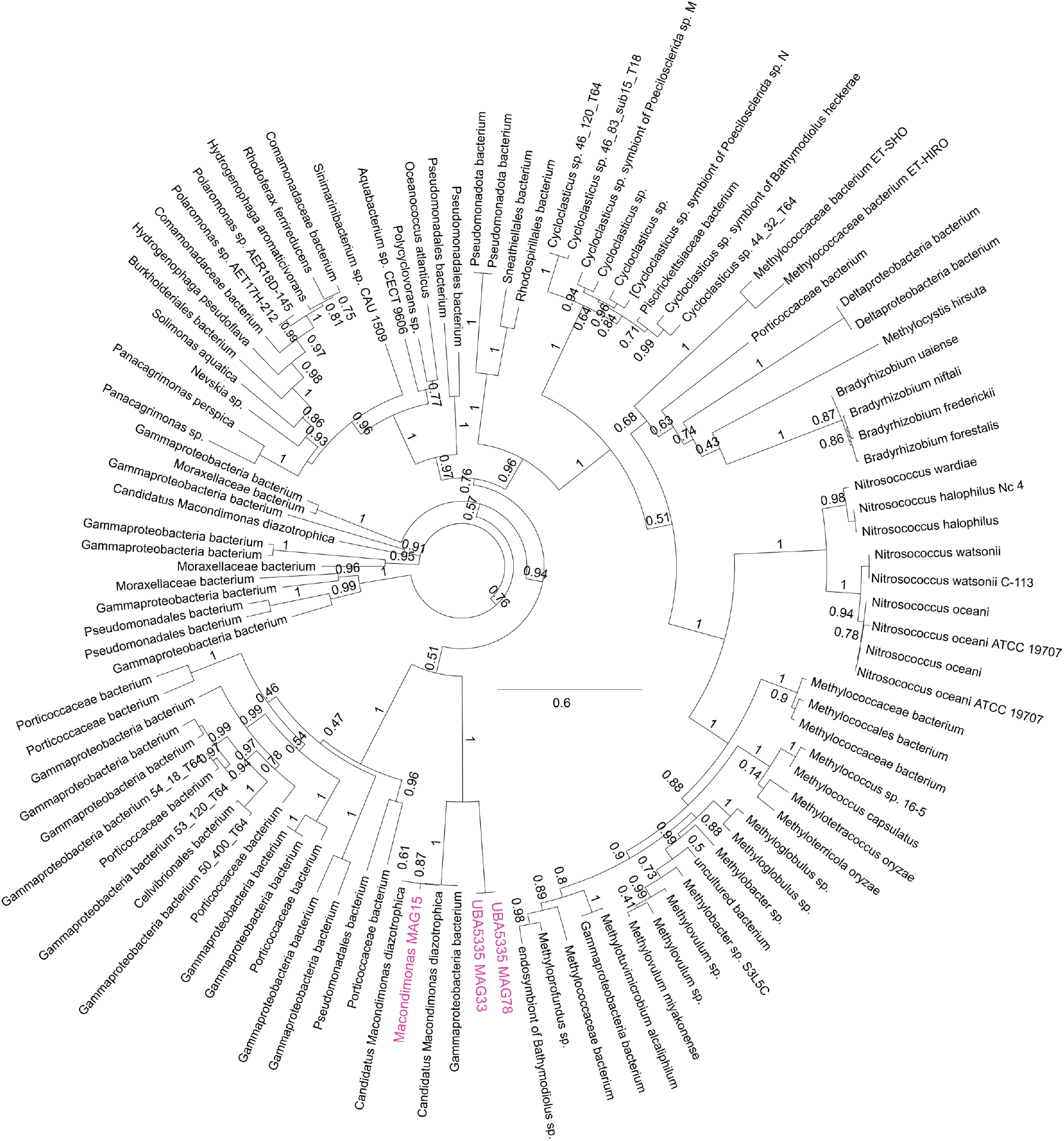
Phylogeny of PhmA proteins in alkane-oxidizing UBA5335, including *Candidatus Macondimonas*, from tar pollution (purple), and related PhmA/PmoA sequences from methane and short-chain alkane oxidizers. Branch bootstrap values are shown, the scale represents the number of substitutions per site.

**Figure S12:**
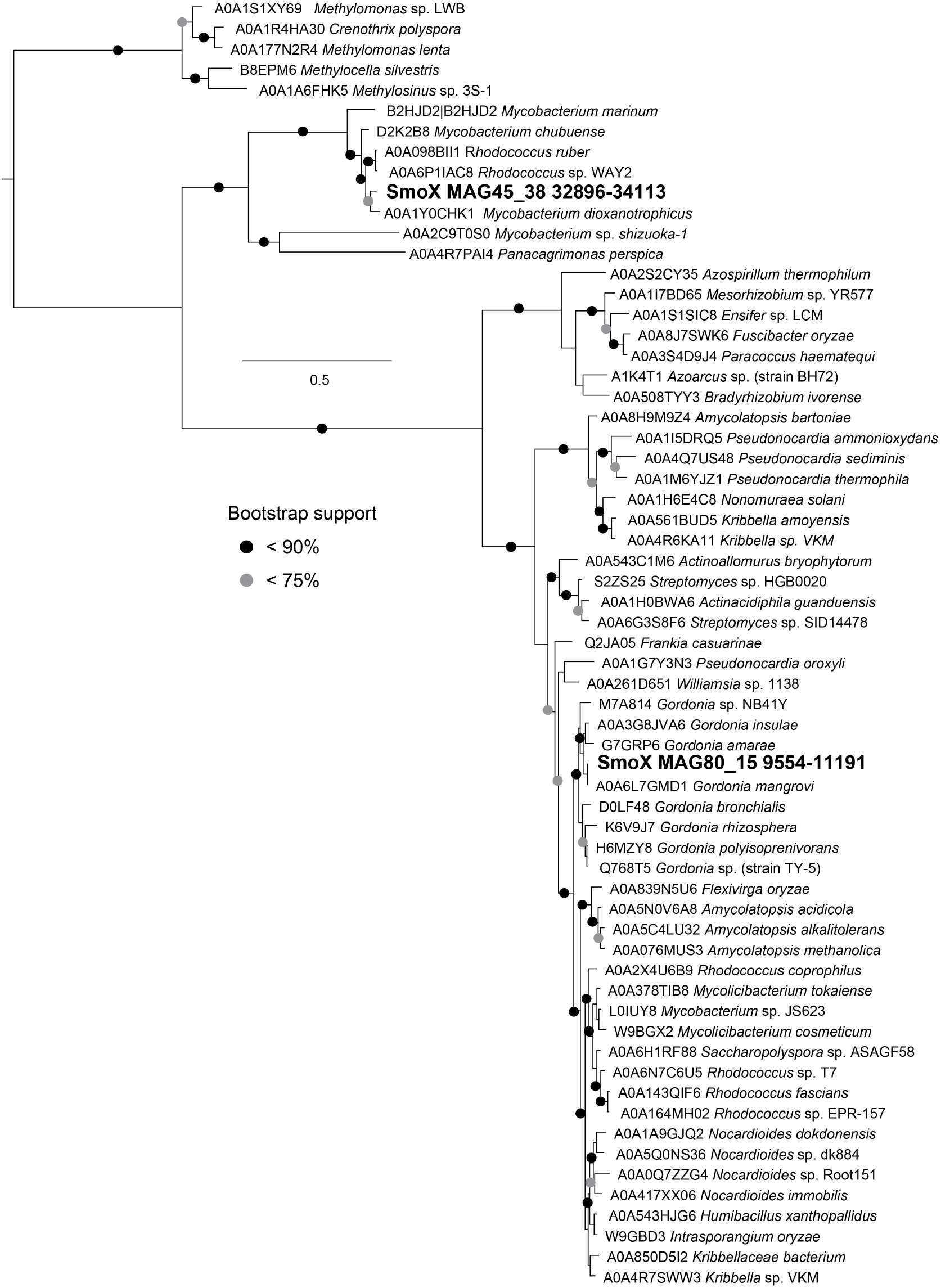
Phylogeny of the soluble methane/propane 2-monooxygenase-like hydroxylase component large subunit SmoX proteins in mycobacteria and related sequenced from UniProt (FastTree, accession numbers shown). Sequences from mycobacterial MAGs 45 and 80 are highlighted. Branch bootstrap values are shown, and the scale represents the number of substitutions per site.

**Figure S13:**
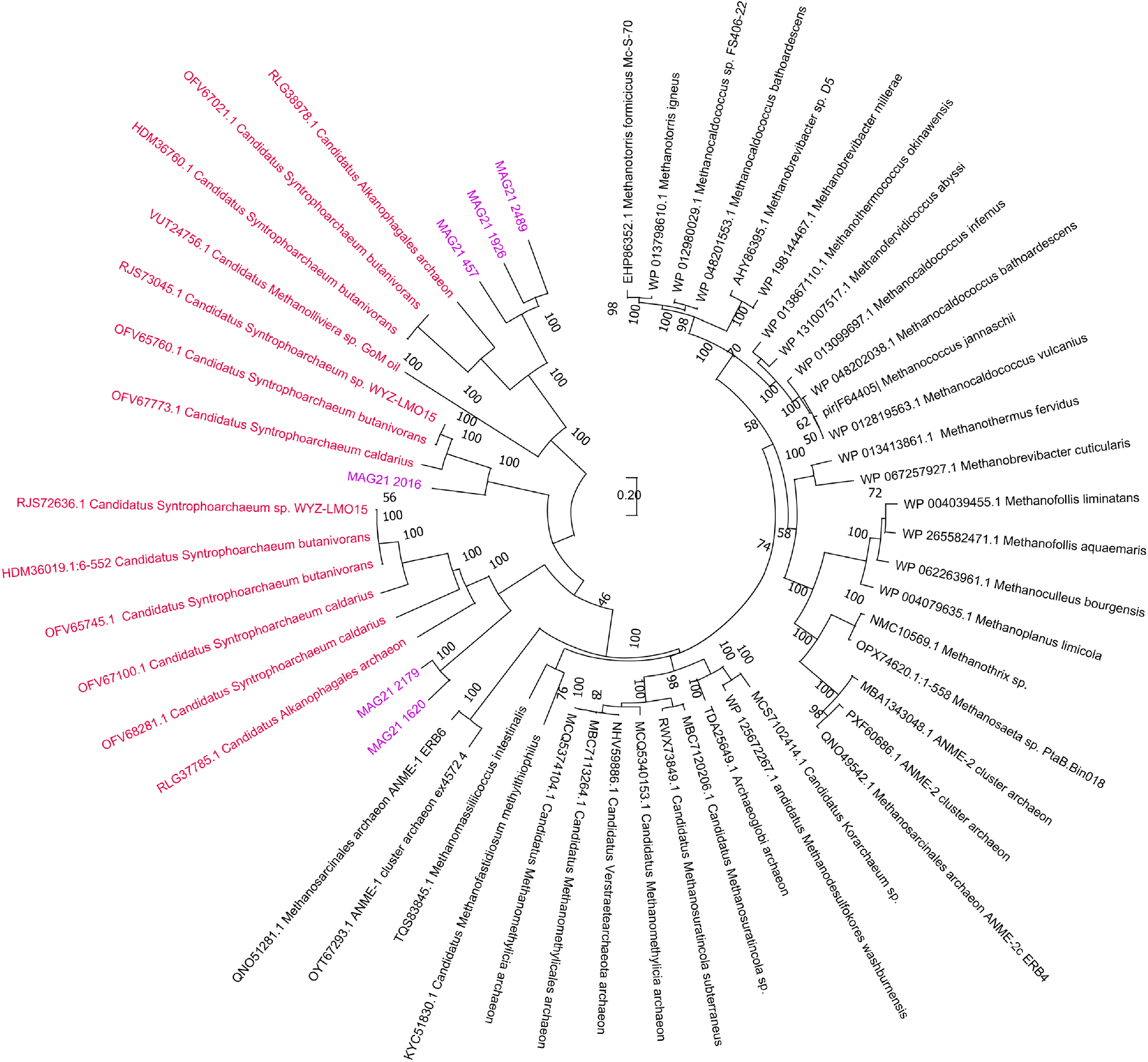
Phylogeny of AcrA/McrA proteins in alkane-oxidizing Syntropharchaeales from tar pollution (purple), the related Syntropharchaeales (red) and methanotrophs and methanogens (unrooted tree, LG+G+I model, MEGA11). Branch bootstrap values are shown, the scale represents the number of substitutions per site. MAG25 also encoded the AcrA/McrA proteins, however, only partial sequences were found in the fragmented assemblies.

**Figure S14:**
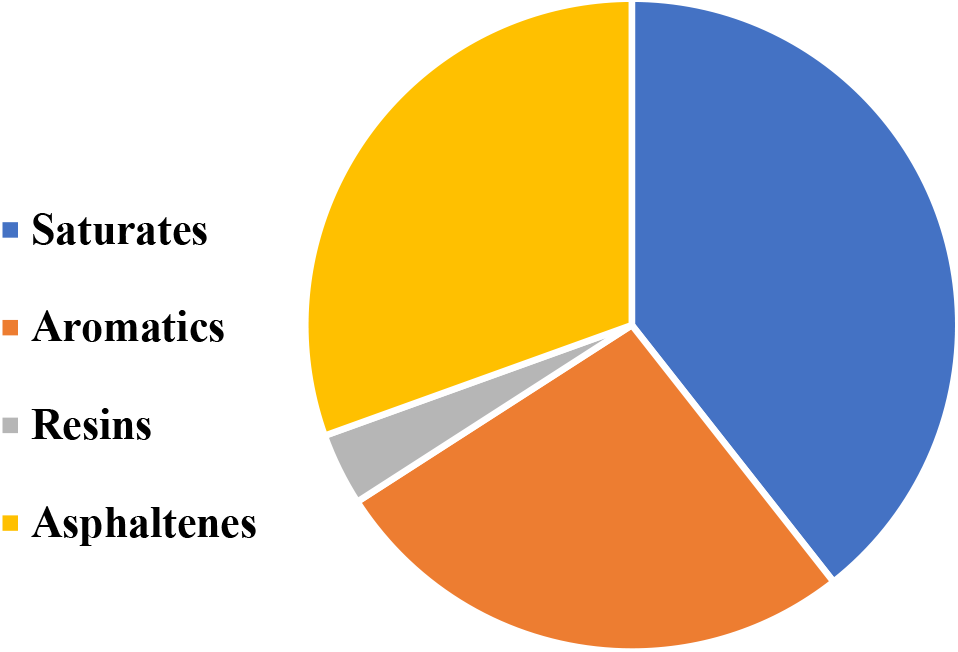
The distribution of the saturates (39.4%), aromatic (26.5%), resins (3.6%) and asphaltenes (30.5%) fractions in the BY2 oil sample.

**Figure S15:**
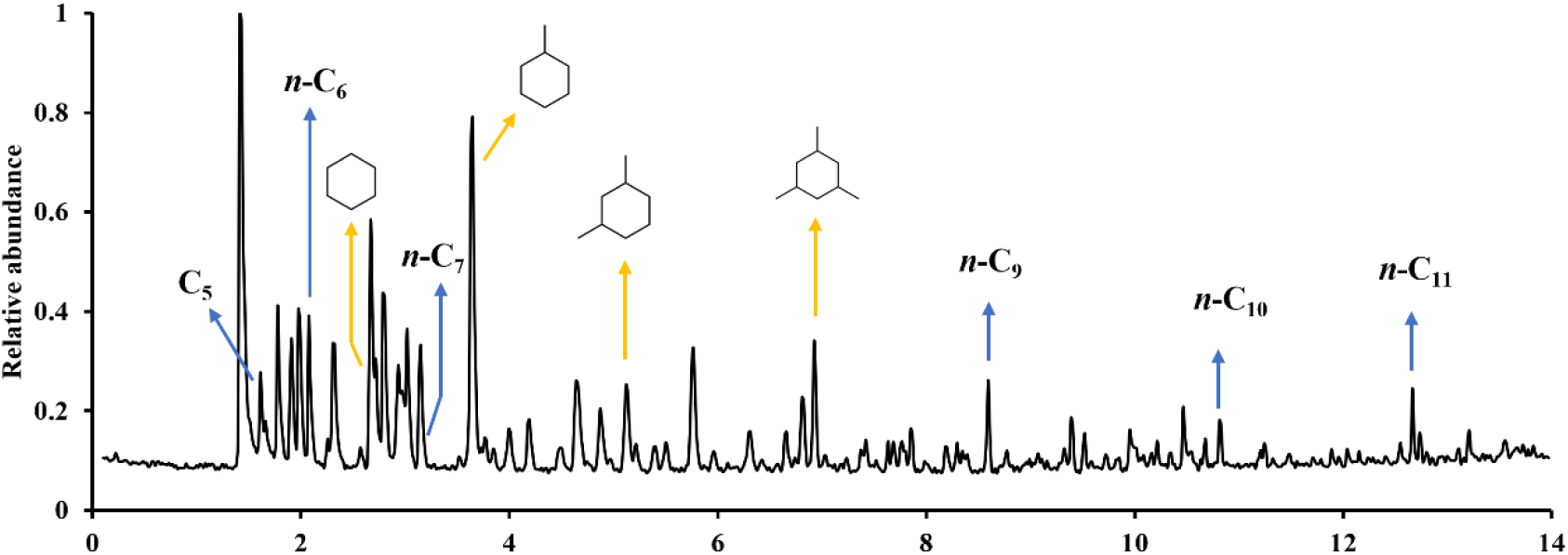
Total ion current (TIC) GC-MS chromatogram of the head-space of BY2 oil sample.

**Figure S16:**
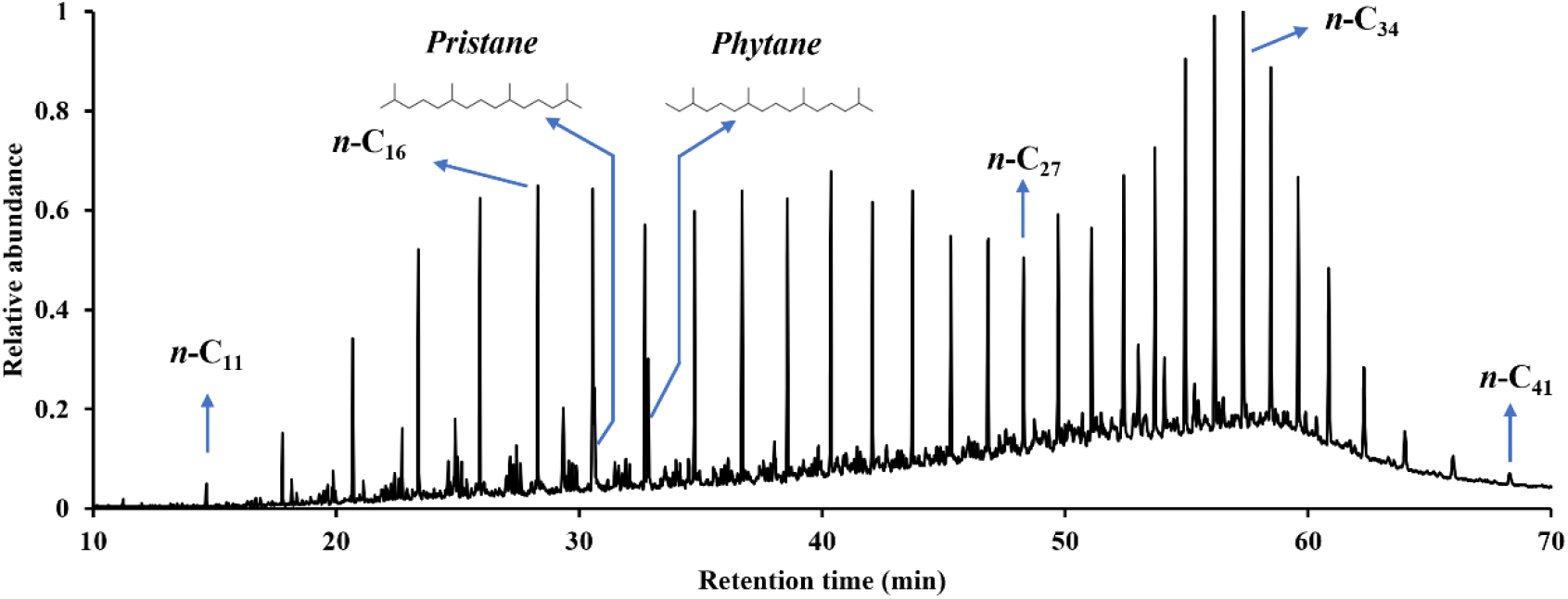
Total ion current (TIC) GC-MS chromatogram showing the n-alkanes distribution, pristane and phytane in the saturate fraction of BY2 oil sample.

**Figure S17:**
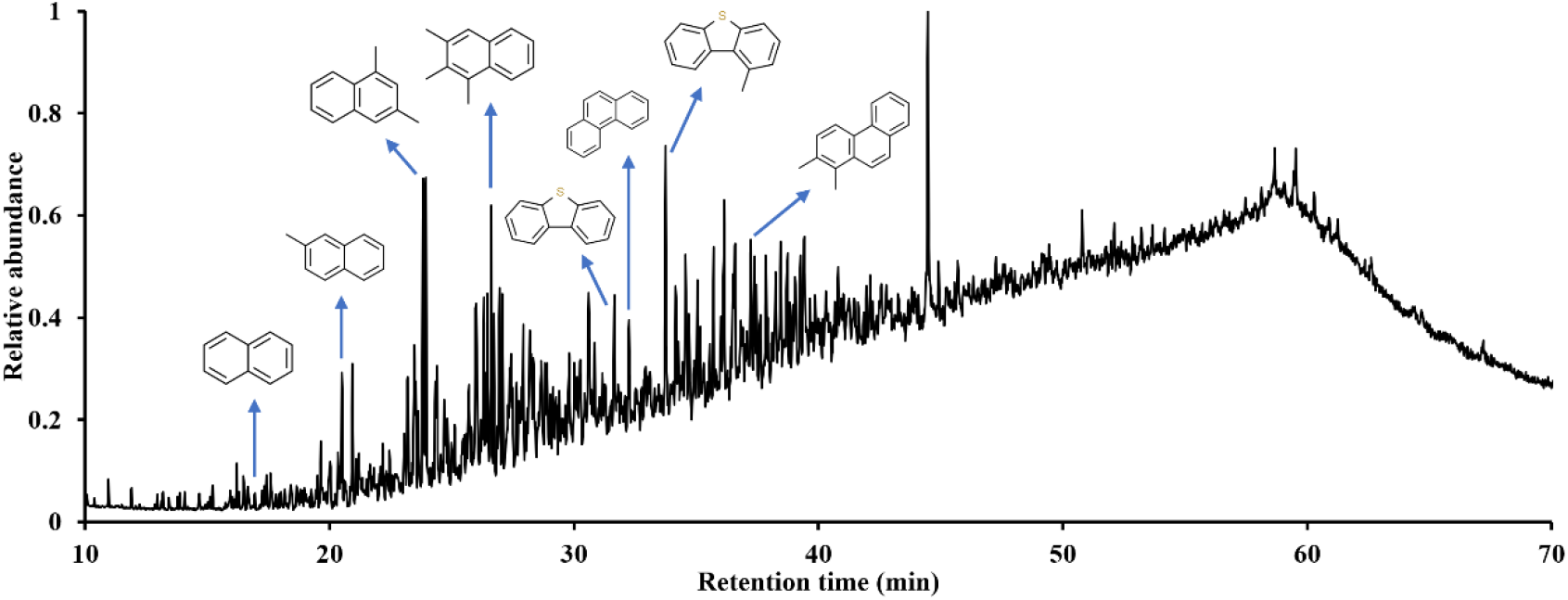
Total ion current (TIC) GC-MS chromatogram showing the distribution of various aromatic compounds in the aromatic fraction of BY2 oil sample.

